# Human USP18 protects diverse cancer lineages from Type I Interferon independently of its canonical catalytic function

**DOI:** 10.1101/2023.03.23.533924

**Authors:** Veronica Jové, Heather Wheeler, Chiachin Wilson Lee, David R. Healy, Kymberly Levine, Erik C. Ralph, Bing Yang, Anand Giddabasappa, Paula Loria, Masaya Yamaguchi, Agustin Casimiro-Garcia, Benedikt M. Kessler, Adán Pinto-Fernández, Véronique Frattini, Paul D. Wes, Feng Wang

**Author notes:** Co-senior authors (Véronique Frattini, Paul D. Wes, Feng Wang).

## Abstract

Precise temporal regulation of Type I interferon signaling is imperative to effectively fight infections and cancerous cells without triggering autoimmunity. The key negative regulator of Type I interferon signaling is ubiquitin-specific protease 18 (USP18). USP18 cleaves interferon-inducible ubiquitin-like modifications through its canonical catalytic function and directly inhibits interferon receptor signaling through its scaffold role. *USP18* loss-of-function dramatically impacts autoimmune disease, viral susceptibility, and cancer cell survival. However, the relative contribution of catalytic versus scaffold function is unresolved and must be determined to design effective therapeutics targeting USP18. To precisely delineate individual contribution, we evaluated the functional impact of single amino acid mutations that disrupt catalytic or scaffold activity. Here we demonstrate catalytic activity does not contribute to cell autonomous Type I interferon sensitivity across multiple cancer cell lineages. Furthermore, introducing a patient-derived mutation that disrupts scaffold function is sufficient to inhibit cancer growth. These findings establish a fundamental mechanistic basis for USP18 therapeutic design across diseases.

**OVERVIEW:**

- USP18 is the key negative regulator of Type I interferon signaling in humans, mediating autoimmune disease, viral susceptibility, and cancer cell survival.
- USP18 cleaves interferon-inducible ubiquitin-like modifications through its canonical catalytic function and attenuates interferon receptor signaling through its scaffold role.
- Delineating the contribution of each function is critical to resolve the mechanistic basis of interferon regulation and enable the development of therapeutics targeting USP18.
- We demonstrate that cell intrinsic interferon sensitivity is not mediated by loss of catalytic activity. However, disruption of scaffold function by a patient-specific mutation inhibits cancer cell growth.
- Furthermore, we discovered that canonical catalytic activity is surprisingly inefficient in human cells.
- These results clarify a fundamental mechanism of immune regulation and cancer cell survival in humans.

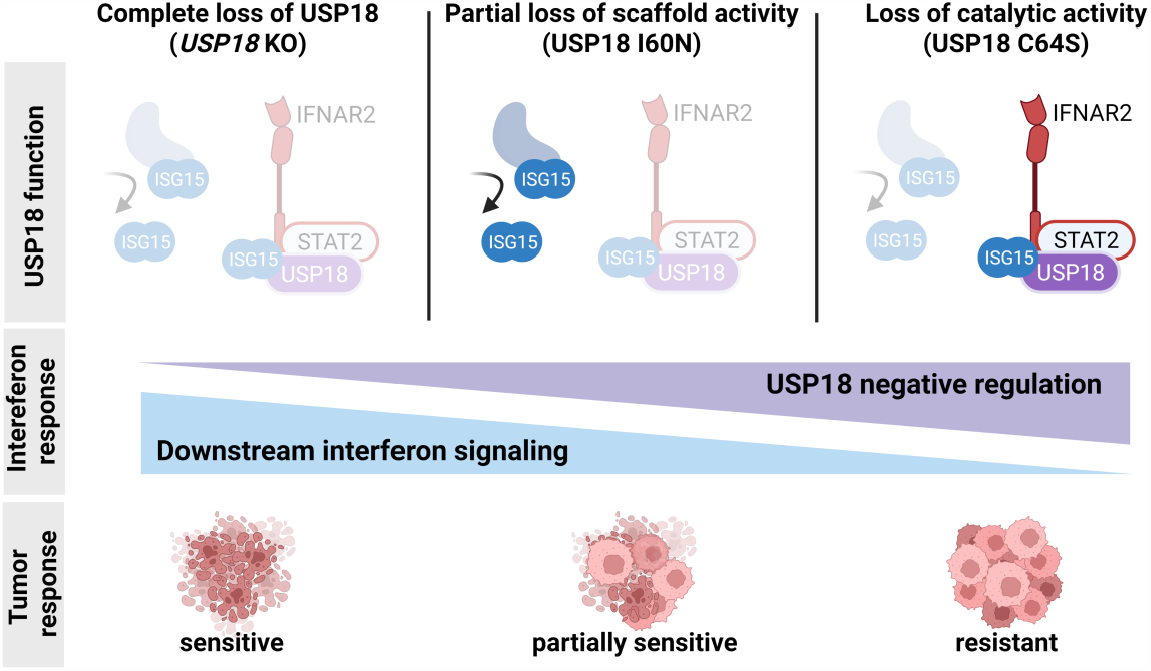

## INTRODUCTION

Type I interferon (IFN) signaling is essential for antiviral defense and regulation of cellular immunity (McNab et al., 2015; Snell et al., 2017). Engagement of the Type I IFN receptor, composed of IFNAR1 and IFNAR2, activates the JAK/STAT signaling cascade, resulting in transcription of interferon stimulated genes (ISGs). ISGs contribute to multiple pro-inflammatory pathways, including antigen processing and presentation, secretion of immunomodulatory chemokines and cytokines, and ultimately, induction of cell autonomous growth arrest and death (Bekisz et al., 2010; Gessani et al., 2014; Schneider et al., 2014). This coordinated inflammatory response eliminates infected and neoplastic cells and can be harnessed therapeutically to promote anti-tumor immunity and tumor intrinsic growth arrest or death (Boukhaled et al., 2021; Cheon et al., 2014; Zitvogel et al., 2015). If left unchecked, however, excessive pro-inflammatory signaling can damage healthy tissue, resulting in auto-immunity (Crow and Stetson, 2022; Taft and Bogunovic, 2018). To prevent excessive inflammation, Type I IFN signaling concomitantly induces negative feedback mediators such as USP18 and PD-L1, which are expressed at undetectable or low levels under basal conditions (Barber et al., 2006; Francois-Newton et al., 2011; Kang et al., 2001).

The central role of USP18 in IFN negative regulation is supported by human genetics (Alsohime et al., 2020; Martin-Fernandez et al., 2022; Meuwissen et al., 2016; Zhang et al., 2015). For instance, *USP18* loss-of-function results in a severe interferonopathy called pseudo-TORCH syndrome 2 that limits lifespan to infancy (Alsohime et al., 2020; Meuwissen et al., 2016). Affected individuals can exhibit microcephaly and features of severe systemic infection in the absence of a detectable infectious agent (Alsohime et al., 2020; Meuwissen et al., 2016). USP18 is thought to negatively regulate Type I IFN signaling by directly binding to the IFN receptor subunit, IFNAR2, resulting in attenuated signaling (Arimoto et al., 2017; Malakhova et al., 2003). In this scaffold role, USP18 is recruited to IFNAR2 with STAT2 and ISG15, effectively competing with JAK1 to bind IFNAR2, thereby preventing downstream signaling events such as STAT1 phosphorylation (Arimoto et al., 2017; Malakhova et al., 2003; Vasou et al., 2021). Point mutations in USP18 (I60N) and STAT2 (R148Q and R148W) that disrupt the scaffold function cause a rare form of severe Type I interferonopathy in patients (Duncan et al., 2019; Gruber et al., 2020; Martin-Fernandez et al., 2022).

In addition to scaffold function, USP18 possesses proteolytic catalytic activity to cleave the interferon-inducible ubiquitin-like protein, ISG15, from ISG15-conjugated (ISGylated) proteins (Basters et al., 2014; Malakhov et al., 2002). Like USP18, ISG15 and the enzymatic cascade required for ISGylation (E1 activating enzyme, UBA7; E2 conjugating enzyme, UBCH8; E3 ligase HERC5) are ISGs and therefore, tightly regulated upon IFNAR engagement (Jimenez Fernandez et al., 2019). Since newly synthesized proteins are ISGylated co-translationally, ISGs are themselves ISGylated (Durfee et al., 2010). Newly synthesized bacterial and viral proteins are also ISGylated if Type I IFN signaling is induced by infection (Perng and Lenschow, 2018; Tang et al., 2010; Zhang et al., 2019; Zhao et al., 2010).

The functional consequences of ISGylation upon IFNAR engagement are not fully understood but have been best characterized in the context of viral infection. ISGylation of viral proteins is believed to block viral replication, resulting in decreased infectivity (Perng and Lenschow, 2018; Tang et al., 2010; Zhao et al., 2010). While mice with impaired USP18 deISGylase activity have increased ISGylation and increased viral protection, mice lacking the E1 enzyme required for ISGylation, UBA7, exhibit decreased ISGylation and decreased viral protection (Giannakopoulos et al., 2009; Ketscher et al., 2015; Lai et al., 2009). Furthermore, viruses like SARS-CoV-2 have evolved deISGylases that efficiently cleave ISG15 from viral and endogenous proteins, which may serve as an immune evasion mechanism (Freitas et al., 2020; Munnur et al., 2021; Perng and Lenschow, 2018).

In the absence of viral infection, it is less clear whether USP18 deISGylase activity can negatively regulate IFN signaling. The impact of ISGylation on endogenous proteins, including ISGs, has remained more elusive, especially in humans. Previous reports have suggested that ISGylation can alter protein stability by competing for ubiquitin-mediated degradation or protein function by modifying protein-protein interactions (Desai et al., 2006; Fan et al., 2015; Perng and Lenschow, 2018). Therefore, it has been proposed that USP18 catalytic activity can negatively regulate IFN signaling by removing ISGylation from pro-inflammatory ISGs, which in turn, reduces their levels (Malakhov et al., 2003; Okumura et al., 2013; Perng and Lenschow, 2018; Pinto-Fernandez et al., 2020). Our understanding of deISGylase activity has been hindered because *USP18* loss-of-function studies cannot disambiguate the contribution of deISGylase and scaffold activity. USP18 scaffold function acts upstream to directly regulate the entire IFN signaling cascade, which includes expression of USP18, ISG15, and the enzymatic cascade required for ISGylation. Since loss of scaffold function is predicted to increase ISG expression and ISGylation, changes correlated with increased ISGylation upon USP18 loss cannot be directly attributed to loss of catalytic function (Figure S1).

Given the crucial role for USP18 in immune regulation, it is of great interest to determine the mechanism by which it functions. Genetic ablation of *USP18* in human cancer cells confers sensitivity to Type I IFN treatment, associated with an accumulation of ISGylated proteins, increased ISG expression, and cell autonomous growth inhibition and death (Arimoto et al., 2023; Pinto-Fernandez et al., 2020). However, the relative contribution of USP18 deISGylase activity versus scaffold function to cancer survival remains unknown. Published reports have implicated catalytic function because modulation of USP18 increased ISGylation of a wide range of proteins, including ISGs and tumor suppressors that can regulate tumor growth (Guo et al., 2012; Mustachio et al., 2017a; Mustachio et al., 2017b; Park et al., 2016; Pinto-Fernandez et al., 2020). For example, increased ISGylation of ISGs and the tumor suppressor p53 is hypothesized to increase tumor intrinsic IFN signaling or p53-dependent transcription, respectively, resulting in decreased tumor growth (Park et al., 2016). Nevertheless, a causal relationship between deISGylase activity and cancer survival has not been established. To determine the relative contribution of scaffold and catalytic activity, it is necessary to develop tools that selectively impair either function and to evaluate multiple cancer lineages.

We investigated the impact of USP18 deISGylase activity across a broad range of human tumors by introducing a de-ISGylation mutation into the endogenous *USP18* locus. Complete loss of USP18 confers IFN sensitivity to multiple cancer cell lineages, yet strikingly, none are sensitive to loss of deISGylase activity. Furthermore, a patient-derived mutation that disrupts IFNAR2 binding, I60N, is sufficient to confer IFN sensitivity. Collectively these findings demonstrate that deISGylase activity does not repress IFNAR signaling or mediate tumor intrinsic IFN sensitivity. Insight into this key negative regulator of IFN signaling facilitates the development of therapeutics for cancer immunotherapy, acute and chronic viral infection, and autoinflammatory disorders.

## RESULTS

### *USP18* loss confers IFN sensitivity across diverse cancer lineages

Previous work demonstrated that genetic ablation of *USP18* in leukemia-derived HAP1 and colorectal HCT116 cancer cells confers sensitivity to IFN treatment (Pinto-Fernandez et al., 2020). IFN sensitivity was associated with accumulation of ISGylated proteins, increased ISG expression, and ultimately cell growth inhibition and death (Pinto-Fernandez et al., 2020). To determine if *USP18* loss confers IFN sensitivity to human cancer cell lines from diverse lineages, we applied the CRISPR LAPSE method developed at Pfizer (Clustered Regularly Interspaced Short Palindromic Repeats Longitudinal Assay Profiling of Specific Edits, Tuladhar and Oyer, in preparation) (Figure 1A). Using this approach, *USP18* knock-out (KO) pools were generated and divided into three sub-pools. One sub-pool was sequenced for initial KO efficiency, another was passaged under the presence of IFN, and a third was passaged under control conditions. After passaging, *USP18* was sequenced, and the proportion of KO to wild-type (WT) alleles was compared to the initial time point and control conditions, thereby indicating whether *USP18* KO resulted in IFN sensitivity (Figure 1A). Consistent with prior data collected from *USP18* KO HAP1 and HCT116 cells, KO allelic frequency was depleted compared to WT after IFN treatment in HAP1 and HCT116 LAPSE pools (Figure 1B). Furthermore, we observed a decrease in *USP18* KO allelic frequency after 2 weeks of IFN treatment in HT-29 (colorectal), A549 and HCC366 (lung), and HCC1143 (breast) LAPSE pools (Figure 1B). Thus, genetic ablation of *USP18* confers IFN sensitivity to multiple cancer cell lines, including those derived from hematopoietic, colorectal, lung, and breast lineages.

**Figure 1:**
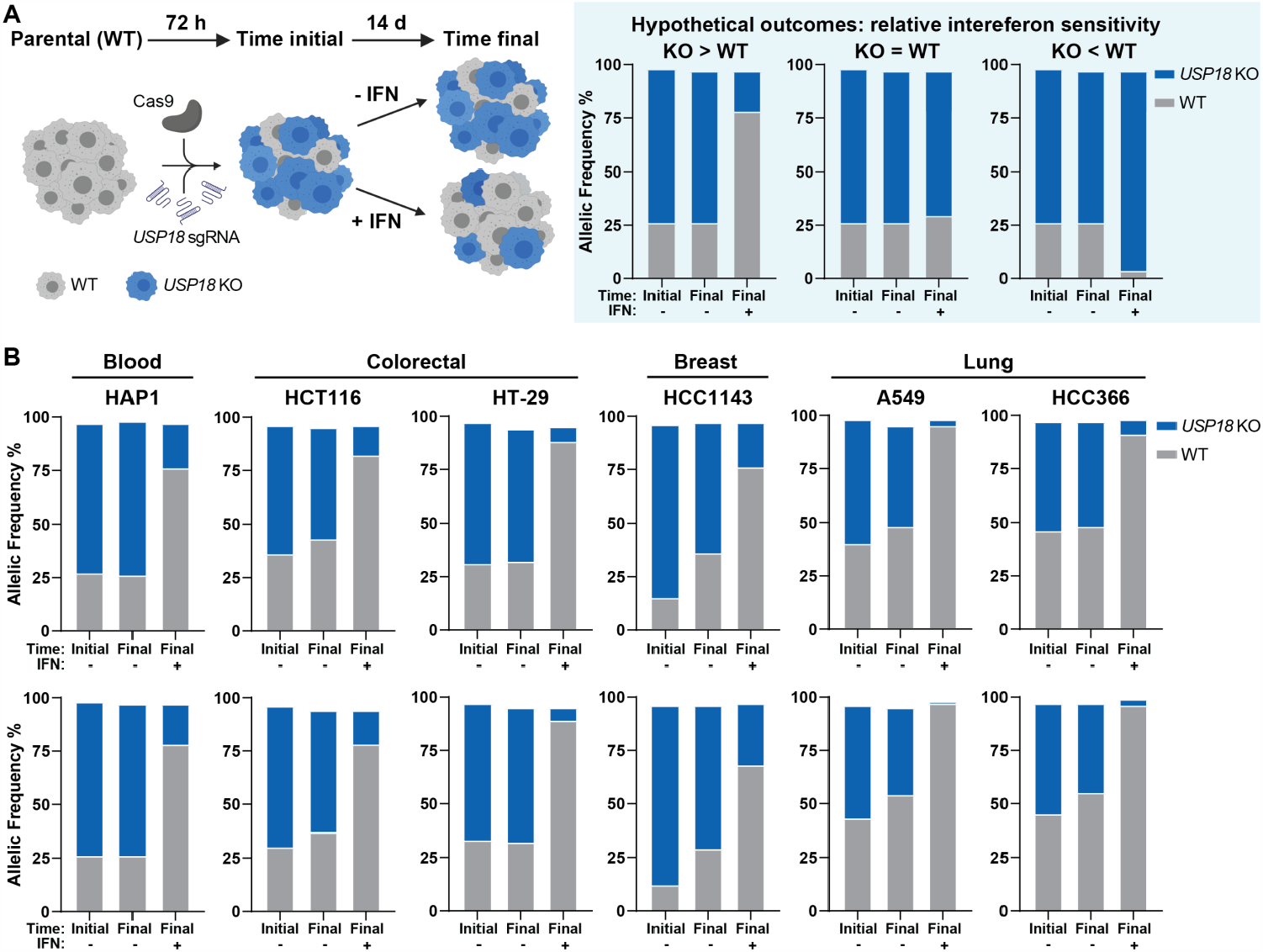
*USP18* loss confers IFN sensitivity across diverse cancer lineages. Parental human cancer cell lines were electroporated with sgRNA targeted to *USP18* and allelic frequency was determined by sequencing samples 72 h post-electroporation (time initial) and after 2 additional weeks of passaging cells (time final) in the presence (+) or absence (−) of 1000 U/mL IFN-α. After 2 weeks of passaging cells, the relative ratio of KO to WT alleles at time final +IFN vs time final -IFN indicates whether *USP18* KO results in IFN sensitivity. (A) Schematic of experimental design (left, created with BioRender.com) and hypothetical outcomes (right) from the CRISPR LAPSE method. (B) KO indicates frame-shift mutation or in-frame mutation ≥ 21 bp; WT indicates no mutation or in-frame mutation < 21 bp. Two independent biological replicates (top and bottom rows) were electroporated, passaged, and sequenced in parallel.

### *Usp18* loss in CT26 tumors promotes tumor growth arrest in vivo

To determine whether *Usp18* loss can inhibit tumor growth in vivo, we genetically ablated *Usp18* in CT26 mouse colorectal cancer cells (*Usp18*^*−/ −*^ CT26). As a control, *Rosa26* was genetically ablated in CT26 cells (*Rosa26*^*−/−*^ CT26). *Usp18*^*−/−*^ or *Rosa26*^*−/ −*^ CT26 tumors were subcutaneously injected in wild-type, immunocompetent BALB/c mice. Tumor volume was significantly reduced in BALB/c mice injected with *Usp18*^*-/-*^ CT26 tumors as compared to *Rosa26*^*−/ −*^ CT26 controls (Figure 2). Therefore, *Usp18* loss in CT26 tumors inhibits tumor growth in vivo.

**Figure 2:**
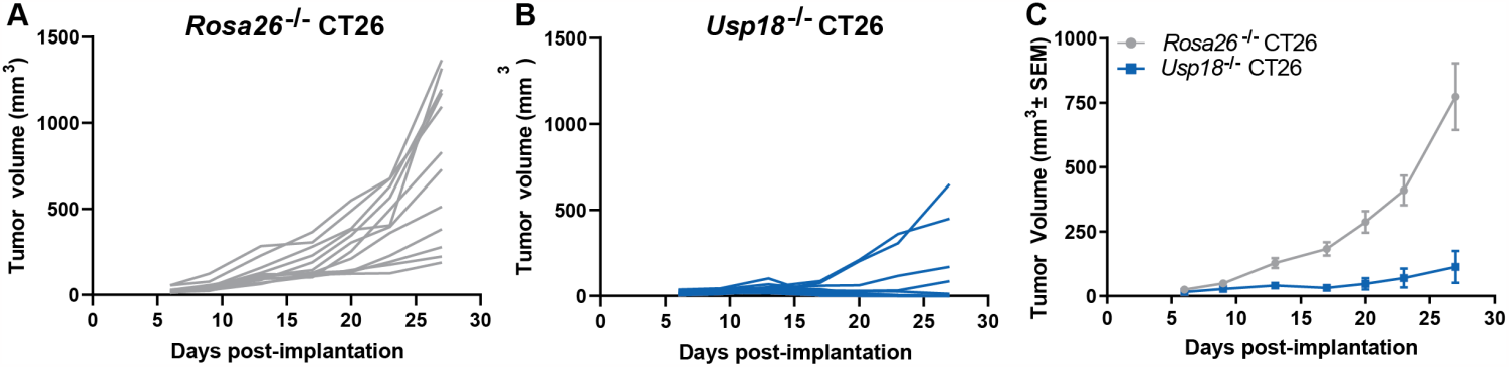
*Usp18* loss in CT26 tumors inhibits tumor growth in vivo. Tumor growth curves of (A,C) control *Rosa26*-/- and (B,C) *Usp18*-/- CT26 tumors injected into wild-type BALB/c mice. Data are represented as (A,B) individual animals and (C) mean c, n = 12 animals per genotype. Results from nonlinear regression analysis presented in Data Table 1 (slopes significantly different fo r each data set, p < 0.0001).

### C64S disrupts USP18 catalytic function

To determine whether loss of USP18 catalytic or scaffold functions, or both, contributed to IFN sensitivity, we introduced single amino acid substitutions predicted to specifically impair each function. Mutation of the catalytic cysteine, C64, to serine (C64S) is predicted to disrupt the ability of USP18 to cleave ISG15 from conjugated proteins (deISGylase activity) while minimizing impact on tertiary protein structure. There is only a single atom change in which the hydroxyl group (−OH) of serine replaces the thiol group (−SH) of cysteine (Corey and Craik, 1992; Craik et al., 1987; Ferreira et al., 2022; Martin-Fernandez et al., 2022; Sprang et al., 1987). A substitution of isoleucine at residue 60 to asparagine (I60N) causes Type I IFN-mediated autoimmunity in patients (Martin-Fernandez et al., 2022; Vasou et al., 2021). The I60N mutation partially inhibits the USP18 scaffold function that represses IFNAR signaling, but spares de-ISGylase activity. While C64S and I60N mutations have been characterized in the literature, the functional impact of these mutations has not been directly compared using recombinant protein or endogenously edited cell lines.

To measure USP18 catalytic activity independently of its scaffold function, we designed two in vitro assays to measure deISGylase activity across USP18 variants. First, we incubated recombinant USP18 with recombinant ISG15 protein conjugated via a peptide bond to a Rhodamine 110 dye (ISG15-Rho110) on its C-terminus. Rho110 is quenched when conjugated to ISG15, and fluorescence increases upon cleavage. WT USP18 showed a slow increase in fluorescence over 80 min when incubated with ISG15-Rho110 (Figure 3A). We hypothesized that inefficient catalytic activity may be related to the high affinity between ISG15 and USP18 in humans (Speer et al., 2016). Therefore, we incubated human USP18 with mouse ISG15-Rho110 substrate, which would presumably have a lower affinity for a non-physiological binding partner. Indeed, the same concentrations of WT USP18 and mouse ISG15-Rho110 reached the maximum limit of fluorescence detection within 10 min (Figure 3A). In contrast to WT, C64S USP18 did not cleave detectable amounts of either human or mouse ISG15-Rho110, indicating that the C64S substitution substantially reduced deISGylase activity (Figure 3A). I60N USP18 activity was comparable to WT USP18 for both human and mouse substrates (Figure 3B). Surprisingly, enzymatic turnover of human ISG15-Rho110 substrate by WT and I60N human USP18 was estimated to be less than a single turnover during the assay time course. In contrast, WT and I60N human USP18 were able to turnover murine ISG15-Rho110, demonstrating the enzyme preparations were kinetically competent with a lower affinity substrate (Speer et al., 2016).

**Figure 3:**
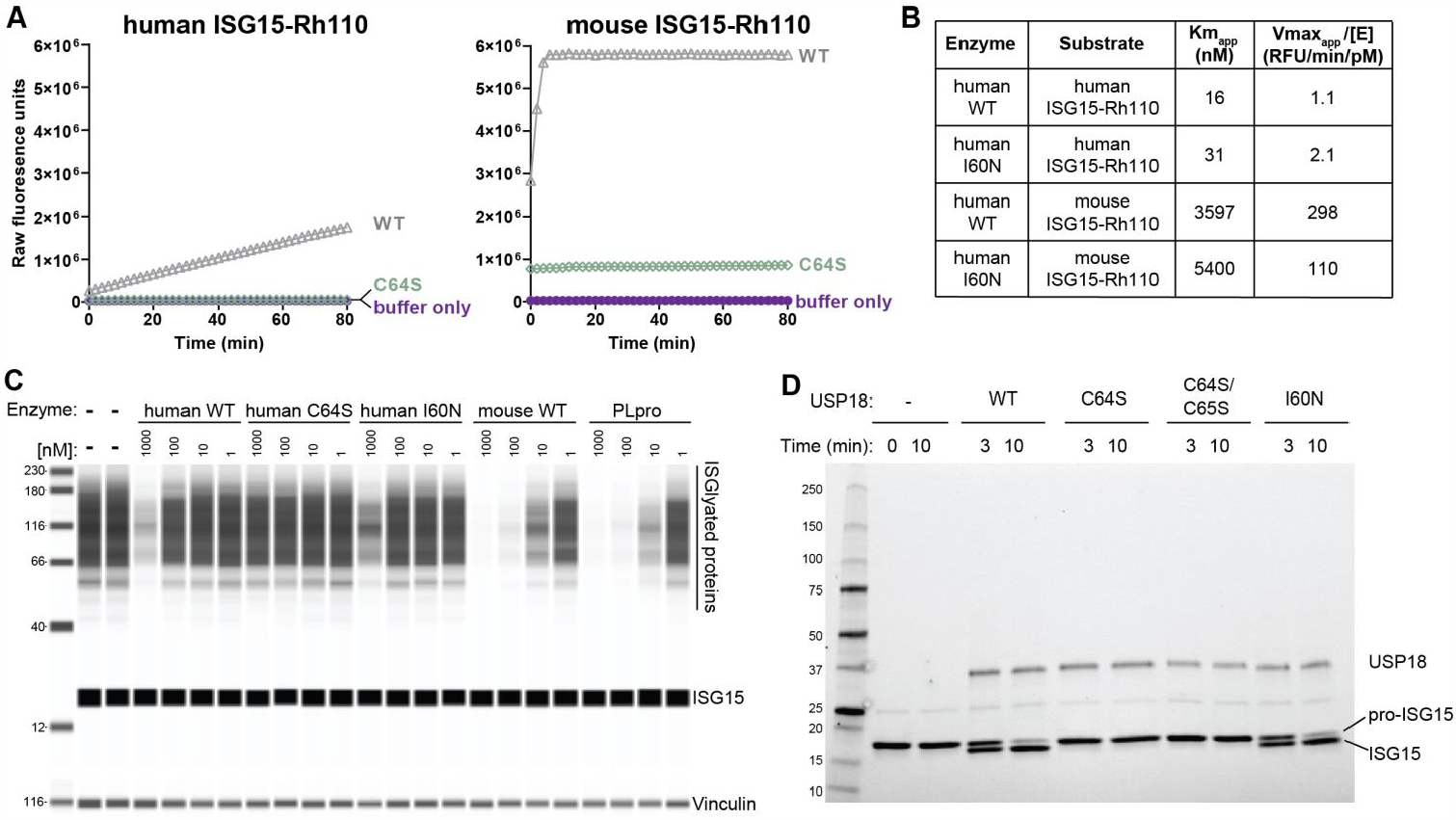
C64S disrupts USP18 catalytic function. (A) Progression curve of 1000 nM human (left) or mouse (right) ISG15-Rho110 cleavage by 10 nM human WT USP18 (WT), human C64S USP18 (C64S), or no enzyme control (buffer only), at RT. (B) Km_apparent_ and Vmax_apparent_ /[Enzyme concentration] determinations for human WT USP18 or human I60N USP18 vs human ISG15-Rho110 or mouse ISG15-Rho110. The reported values are averaged from 2 determinations at different enzyme concentrations and rounded to 2 significant digits. (C) HAP1 *USP18* KO cells were treated for 24 h with IFN-α prior to cell lysis and lysates were incubated with 1, 10, 100, or 1000 nM of indicated recombinant protein for 1 h at RT. Lysates were analyzed by western blot for levels of ISGylated proteins and ISG15 (top) or vinculin (bottom). (D) Cleavage of 5 μM human pro-ISG15 (AA1-165) to mature ISG15 (AA1-157) was assessed by SDS-PAGE after 10 min incubation at 37ºC with 1 μM of recombinant human WT USP18 (WT), human C64S USP18 (C64S), human C64S/C65S USP18 (C64S/C65S), human I60N USP18 (I60N), or no enzyme control (−).

Building on these results, we next assessed deISGylation of endogenously ISGylated proteins by WT, C64S, and I60N USP18. To induce endogenous ISGylation, *USP18* KO HAP1 cells were treated with 1000 U/mL IFN for 24 h. The resulting cell lysates were incubated with WT, C64S, or I60N human USP18 for 1 h and remaining ISGylated proteins were visualized by western blot. As positive controls, mouse WT USP18 and the SARS-CoV-2 deISGylase and deubiquitinase PLpro were run in parallel (Freitas et al., 2020). A decrease in ISGylated proteins was observed upon incuba**tion with ≥ 10 nM of mouse WT USP18 and PLpro** (Figure 3C). Decreased ISGylated substrate levels were also observed upon incubation with 1000 nM WT or I60N USP18 (Figure 3C). Although I60N USP18 was catalytically competent in both in vitro assays, we observed a further decrease in endogenous ISGylated protein levels upon incubation with 1000 nM WT USP18 as compared to incubation with 1000 nM I60N USP18 (Figure 3C). In agreement with the ISG15-Rho110 assay, incubation with C64S did not alter de-ISGylation levels (Figure 3C). Thus, the C64S mutation impairs deISGylation of ISG15-Rho110 and endogenously ISGylated proteins.

Finally, USP18 has peptidase catalytic activity enabling it to cleave pro-ISG15 (AA 1-165) into mature ISG15 (AA 1-157) (Malakhov et al., 2002). We evaluated how USP18 variants impact ISG15 cleavage by incubating 1 μM of WT, C64S, C64S/C65S, or I60N USP18 with 5 μM of recombinant pro-ISG15 for 3 or 10 min. ISG15 cleavage was assessed by SDS-PAGE. Mature ISG15 was detected after incubation with WT and I60N, but not C64S or C64S/C65S, USP18 (Figure 3D). Pro-ISG15 cleavage was not observed in the absence of recombinant USP18 protein. The expected mass for pro- and mature ISG15 protein products was confirmed by mass spectrometry (Figure S2). Together these data demonstrate that the C64S substitution impairs both established catalytic functions of USP18.

### I60N disrupts USP18 scaffold function

Having determined catalytic function with purified proteins, we next introduced these single amino acid mutations into the endogenous *USP18* locus to create C64S or I60N knock-in (KI) cells. In contrast to exogenous constitutive overexpression, regulation by endogenous regulatory elements ensures USP18 expression is induced only in response to IFN and is equivalent between cell lines expressing WT, C64S, and I60N USP18. In its scaffold capacity, USP18 binds to IFNAR2 and attenuates IFNAR signaling, resulting in decreased phosphorylation of STAT1 and STAT2 downstream of IFNAR stimulation. To quantify scaffold activity of USP18 variants, we applied Homogenous Time-Resolved Fluorescence (HTRF) assays and measured the ratio of phosphorylated STAT1 (pSTAT1) to total STAT1 levels. Cells were pre-treated with 1000 U/mL of IFN for 4 h, washed, and rested overnight to induce ISGs, including USP18, STAT2, and ISG15. Cells were then re-stimulated with a pulse of IFN for 15 min, and the levels of pSTAT1 and total STAT1 were measured. *USP18* KO cells demonstrated a complete loss of scaffold function as shown by the increase in pSTAT1/total STAT1 levels compared to WT (Figure 4). While partial scaffold deficits were observed in I60N KI cells, C64S KI cells showed no deficits in scaffold function compared to WT (Figure 4). Surprisingly, C64S KI cells exhibited a modest enhancement of scaffold function (Figure 4A). Since USP18 expression was induced post-IFN stimulation, no differences were observed between WT, *USP18* KO, and KI cells without IFN pre-treatment (Figure 4).

**Figure 4:**
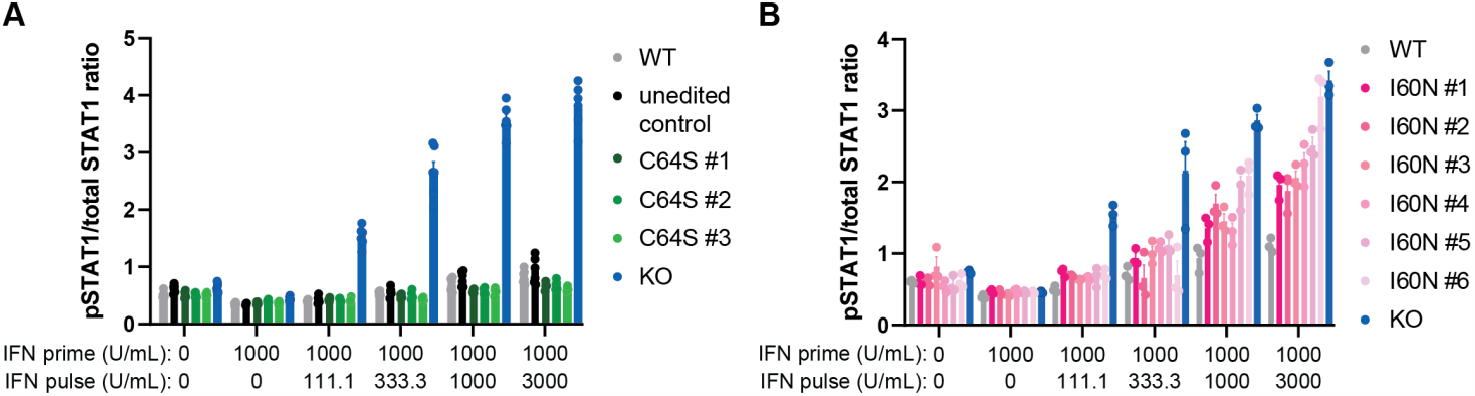
I60N disrupts USP18 scaffold function. (A,B) Ratio of phosphorylated STAT1 (pSTAT1) to total STAT1 levels measured by HTRF in individual HAP1 *USP18* KO and (A) USP18 C64S KI (C64S) or (B) USP18 I60N KI (I60N) clones compared to parental HAP1 cells (WT). Cells were treated with 1000 U/mL IFN-α for 4 h (IFN prime), followed by 24 h rest, and subsequent 15 min IFN-α re-stimulation (IFN pulse) at the indicated concentration prior to cell lysis (mean ± SEM, n = 3 replicates). (A) Unedited control clone underwent same electroporation conditions as C64S KI clones, but no editing was observed at the endogenous *USP18* locus. Results from 2-way ANOVA test with Tukey’s multiple comparisons presented in Data Table 1.

Together these experiments demonstrate that the C64S substitution impairs catalytic function and the I60N substitution partially impairs scaffold function. Therefore, C64S and I60N substitutions are effective tools to interrogate the impact of inhibiting catalytic or scaffold activity, respectively, in cancer cells.

### Disrupting deISGylation is not sufficient to confer IFN sensitivity

It is unknown which functional activities of USP18 contribute to the increased ISG induction, ISGylation, and cell growth inhibition upon IFN treatment of *USP18* KO cells. Thus, we assayed C64S and I60N KI cells to determine if preventing deISGylation is sufficient to phenocopy *USP18* KO cells. Upon 48 h of 1000 U/mL IFN treatment, ISG induction, ISGylated protein levels, and cell growth inhibition were significantly reduced in C64S KI compared to *USP18* KO cells (Figure 5A, 5C, 5D). Although C64S KI and WT cells exhibited similar phenotypes, C64S KI cells had a modest decrease in ISG expression, ISGylation, and cell growth inhibition compared to WT cells (Figure 5A, 5C, 5D). Decreased IFN sensitivity in C64S KI cells may be due to the small enhancement of scaffold function observed in the STAT1 HTRF assay (see Figure 4A) (Vasou et al., 2021). Therefore, preventing deISGylation is not sufficient to phenocopy *USP18* KO cells. I60N KI cells, however, exhibited intermediate levels of ISG expression, ISGylation, and cell growth inhibition compared to *USP18* KO and WT cells (Figure 5B, 5E). These results indicate a correlation between impaired scaffold function and increased IFN sensitivity. However, we cannot formally exclude the possibility that I60N disrupts additional undefined scaffold functions of USP18 beyond IFNAR2 repression (Martin-Fernandez et al., 2022; Vasou et al., 2021). Nonetheless, impairing deISGylase activity of USP18 is not sufficient to confer IFN sensitivity.

**Figure 5:**
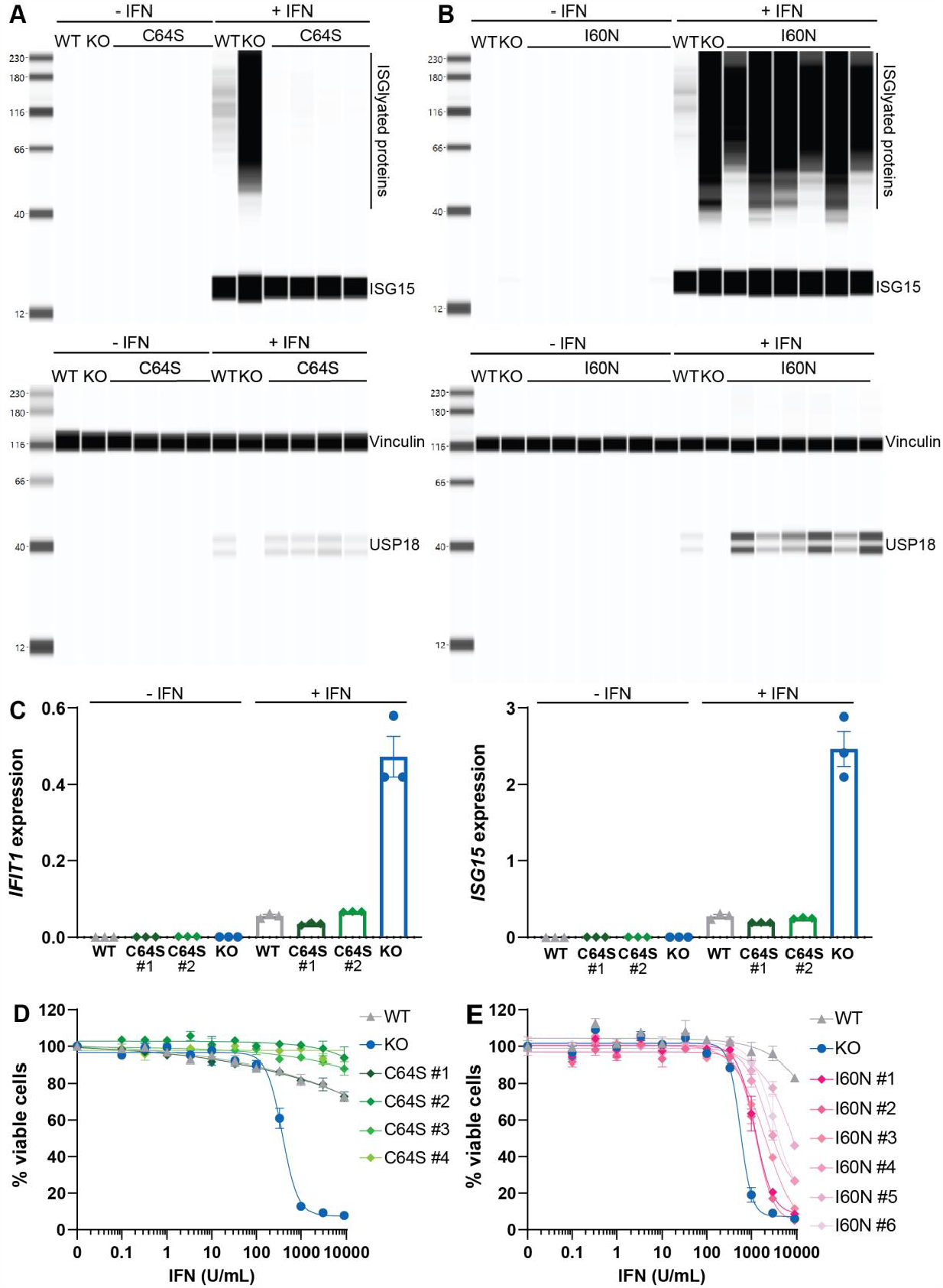
Disrupting deISGylation is not sufficient to confer IFN sensitivity. Individual HAP1 *USP18* KO (KO) and (A,C,D) USP18 C64S KI (C64S) or (B,E) USP18 I60N KI (I60N) clones were compared to parental HAP1 cells (WT). (A,B) Whole cell lysates were analyzed by western blot for levels of ISGylated proteins and ISG15 (top) or vinculin and USP18 (bottom). Cells were treated with 1000 U/mL IFN-β for 24 h prior to cell lysis. (C) Relative mRNA expression of *IFIT1* (left) or *ISG15* (right) to *GAPDH* as measured by qRT-PCR. Cells were treated with 1000 U/mL IFN-β for 48 h prior to cell lysis (mean ± SEM, n = 3 replicates). Results from Kruskal-Wallis test with Dunn’s multiple comparisons presented in Data Table 1. (D,E) Cell viability was measured after 72 h treatment with indicated con-centration of IFN-α. Viability was normalized to 0 U/mL IFN control for each cell line. Each data point denotes mean of n = 2 replicates ± SEM. EC50 determinations for each cell line are presented in Data Table 1.

### Disrupting the ISGylation pathway does not rescue IFN sensitivity

If deISGylase activity does not mediate IFN sensitivity, *USP18* KO cells with defects in the ISGylation pathway should remain sensitive to IFN. To this end, we genetically ablated the E1 ligase required for ISGylation, UBA7 (Ubiquitin-Like Modifier Activating Enzyme 7), in WT and *USP18* KO cells to generate *UBA7* KO and *UBA7/USP18* double KO (*UBA7/USP18* dKO) cells (Figure 6A, 6B). As expected, a substantial decrease in ISGylation was observed in *UBA7* KO and *UBA7/USP18* dKO cells upon 24 h 1000 U/mL IFN treatment (Figure 6A). Furthermore, *UBA7/USP18* dKO cells remained sensitive to IFN and induced ISG expression at levels comparable to *USP18* KO cells (Figure 6C, 6D). No significant differences in ISG induction or IFN sensitivity were observed between *UBA7* KO cells and WT (Figure 6C, 6D). By genetically ablating the E1 ligase required for ISGylation, UBA7, we demonstrate that loss of ISGylation does not rescue IFN sensitivity upon *USP18* KO. These results provide orthogonal evidence that the deISGylase activity of USP18 is dispensable for IFN sensitivity.

**Figure 6:**
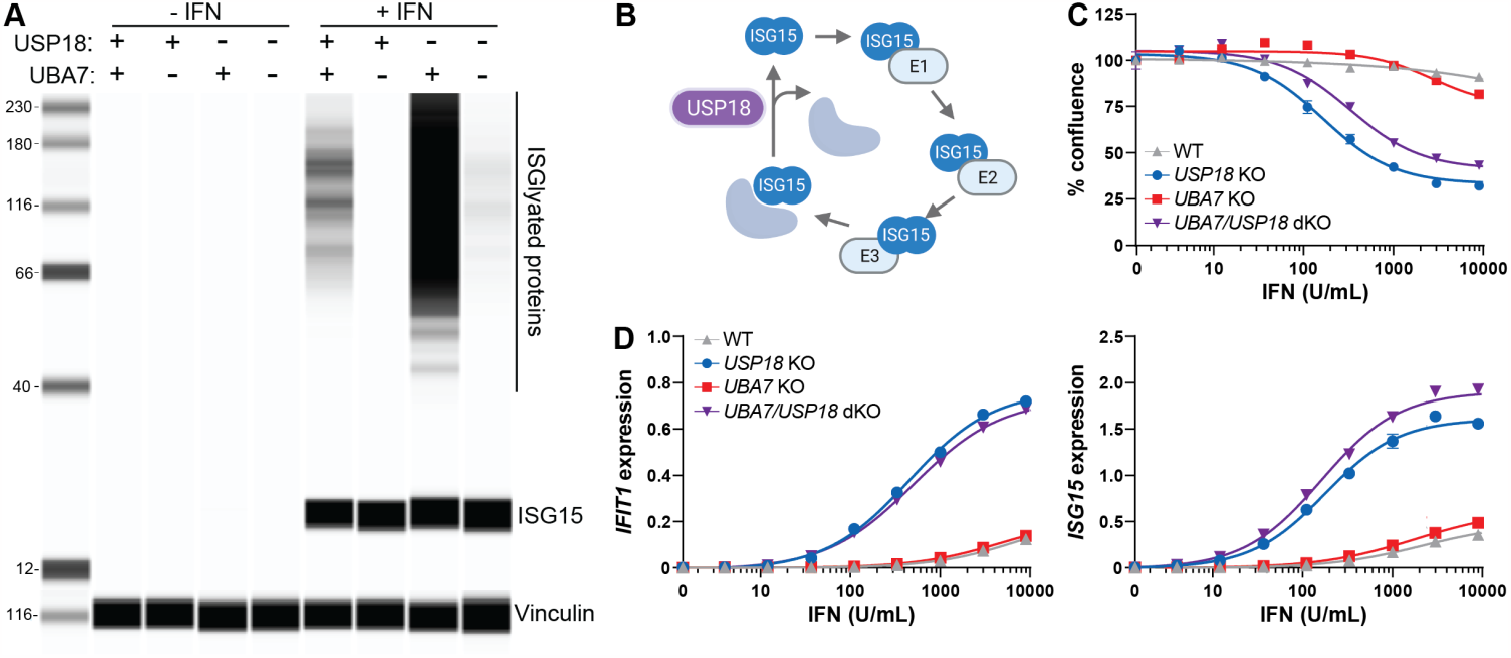
Disrupting the ISGylation pathway does not rescue IFN sensitivity. *UBA7* KO pools and *UBA7/USP18* double KO (dKO) pools were compared to *USP18* KO clone and parental HAP1 cells (WT). (A) Whole cell lysates were analyzed by western blot for levels of ISGylated proteins and ISG15 (top) or vinculin (bottom). Cells were treated with 1000 U/mL IFN-β for 24 h prior to cell lysis. (B) Schematic of the enzymatic cascade required for ISGylation (UBA7, UBCH8, and HERC5) and deISGylation (USP18), created with BioRender.com. (C) Cell viability and (D) relative mRNA expression of *IFIT1*(left) or *ISG15* (right) to *GAPDH* measured by RT-qPCR after 48 h treatment with indicated concentration of IFN-β. Each data point denotes mean of n = 3 replicates ± SEM. Confluence was normalized to 0 U/mL IFN control from each cell line and EC50 determinations for each cell line are presented in Data Table 1.

### DeISGylation does not mediate IFN sensitivity across diverse cancer lineages

To determine if deISGylation is dispensable for IFN sensitivity across multiple cancer cell lines, we performed CRISPR LAPSE experiments with C64S knocked-in to the endogenous *USP18* locus of blood (HAP1), breast (HCC1143, HCC1954, SKBR3, MDA-MB-468), colorectal (HCT116 and

HT-29), and lung (A549, NCI-H1650, HCC366) cancer cells. Initial C64S KI and KO efficiency in pools were measured 72 h after electroporation using genomic DNA PCR prior to 2 weeks ± IFN treatment. Consistent with the observed IFN sensitivity in HAP1 *USP18* KO clones, the frequency of HAP1 *USP18* KO cells decreased upon IFN stimulation in LAPSE experiments (Figure 7A and Figure S3). Although the frequency of WT was relatively stable between HAP1 pools with or without IFN treatment, the frequency of C64S increased upon IFN treatment, in line with the observation that C64S KI cells have a modest decrease in IFN sensitivity compared to WT cells (Figure 7A and Figure S3). The modest enhancement of C64S scaffold function observed upon brief (15 min) IFN pulses in HTRF experiments may cumulatively confer significant protection during prolonged (2 weeks) IFN selective pressure. KO cells were depleted compared to WT upon IFN stimulation across all pools tested except for MDA-MB-468 (Figure 7A and Figure S3). MDA-MB-468 cells contain a single nucleotide polymorphism splice variant in STAT2, which may alter Type I IFN signal transduction and/or the scaffold interaction between USP18 and STAT2 required for IFNAR repression. Importantly, the frequency of C64S cells did not decrease across all pools tested (Figure 7 and Figure S3). In fact, as with HAP1 cells, the frequency of C64S increased in most cancer cell lines treated with IFN as compared to no IFN controls (Figure 7 and Figure S3). Therefore, inhibiting deISGylation is not sufficient to confer IFN sensitivity across multiple cancer cell lines, including those derived from blood, colorectal, breast, and lung lineages (Figure 8).

**Figure 7:**
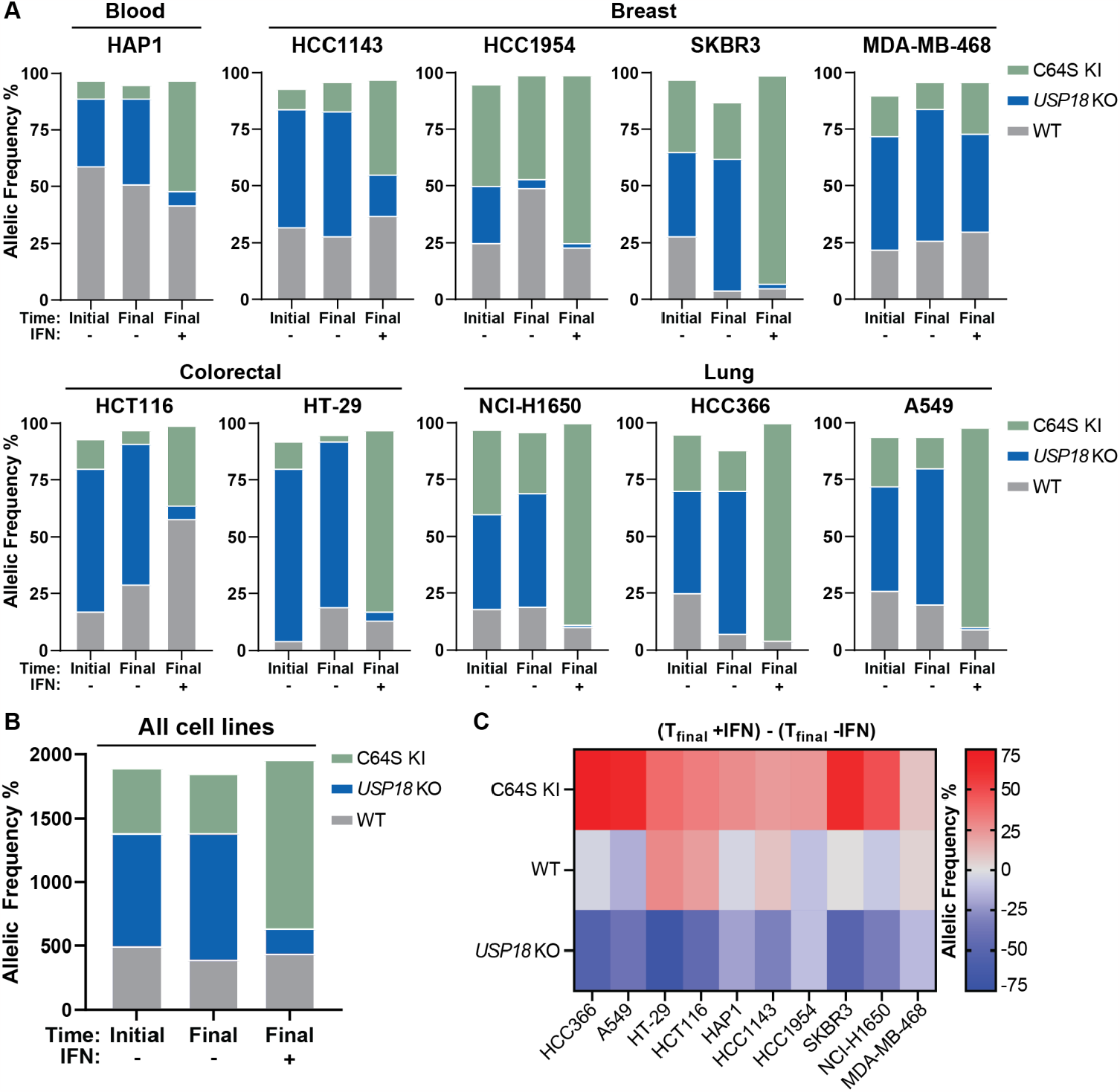
DeISGylation does not mediate IFN sensitivity across diverse cancer lineages. (A) Parental human cancer cell lines were electroporated with C64S donor oligo and sgRNA targeted to *USP18*. Allelic frequency was determined by sequencing samples 72 h post-electroporation (time initial) and after 2 additional weeks of passaging cells (time final) in the presence (+) or absence (-) of 1000 U/mL IFN-α. KO indicates frame-shift mutation or in-frame mutation ≥ 21 bp; WT indicates no mutation or in-frame mutation < 21 bp; KI indicates donor oligo integration at the appropriate position. An independent biological replicate was electroporated, passaged, and sequenced in parallel (Figure S3). (B,C) Summary of data presented in (A) and Figure S3. (B) Cumulative percent allelic frequency for each genotype (C64S KI, *USP18* KO, WT) at the indicated time point, summed across all cell lines and biological replicates. (C) Percent allelic frequency for each genotype at time final -IFN was subtracted from time final +IFN. Positive or negative values indicate an increase or decrease, respectively, in allelic frequency after IFN treatment. Each box denotes the mean of 2 biological replicates.

**Figure 8:**
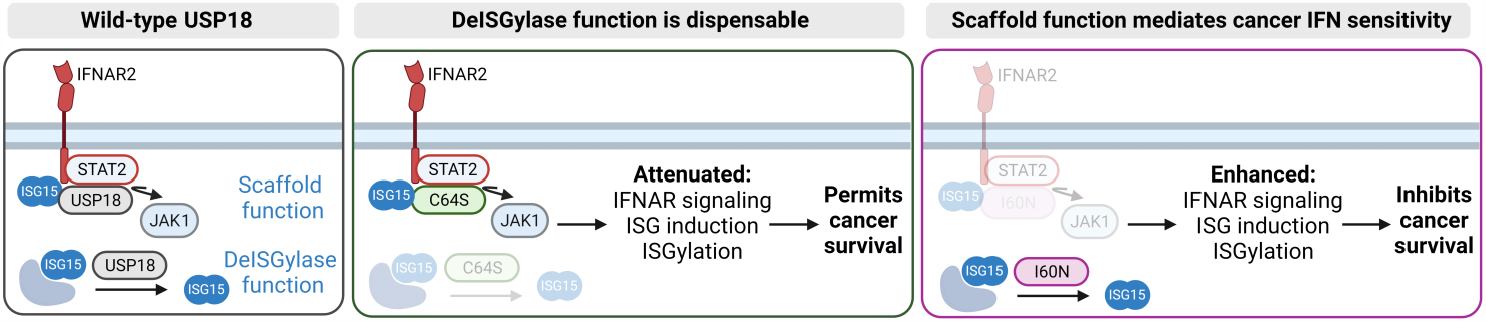
Proposed model for USP18-mediated IFN sensitivity in cancer. Wild-type USP18 negatively regulates Type I IFN responses through its catalytic function to deISGylate ISG15-conjuagated proteins and its scaffold function to repress IFNAR signaling by displacing JAK1. Although wild-type USP18 protects cancer cells from Type I IFN sensitivity, complete loss-of-function mutations in USP18 result in enhanced ISG induction, ISGylation, and cell growth inhibition. Loss of deISGylase activity in USP18 C64S mutants does not confer IFN sensitivity and USP18 C64S mutants exhibit a comparable phenotype to WT. However, partial loss of scaffold function in USP18 I60N mutants results in partial IFN sensitivity and an intermediate phenotype between WT and *USP18* KO. Model created with BioRender.com.

## DISCUSSION

USP18 is a key negative regulator of Type I IFN signaling, yet the relative contribution of catalytic versus scaffold function has not been firmly established. Published reports have implicated either activity, complicating the development of therapeutics targeting USP18 (Guo et al., 2012; Mustachio et al., 2017a; Mustachio et al., 2017b; Park et al., 2016; Pinto-Fernandez et al., 2020; Potu et al., 2010). In this manuscript, we demonstrate that deISGylase activity is not required to suppress Type I IFN signaling in human cancer cells (Figure 8).

Prior work demonstrated that ectopic expression of USP18 C64R/C65R mutants could not rescue IFN sensitivity in HAP1 *USP18* KO cells (Pinto-Fernandez et al., 2020). However, the contribution of USP18 catalytic versus scaffold activity to IFN sensitivity could not be definitively determined in this system because the bulky double arginine substitutions may cause structural changes that impair USP18 scaffold function in addition to catalytic function. For example, cells expressing C64R/C65R exhibited increased ISG expression, suggesting impaired scaffold function (Pinto-Fernandez et al., 2020). Furthermore, we reasoned that overexpression using an exogenous promoter would lack contemporary regulation of other IFN stimulated genes, which may be important to fully recapitulate USP18 function. In this study, we introduced a deISGylation-specific mutation into the endogenous *USP18* locus. These results clearly demonstrate that while a broad range of cancer cell types are sensitive to IFN upon complete USP18 loss, none are sensitive to loss of deISGylase activity.

Our biochemical results indicate that human USP18 inefficiently deISGylates its endogenous substrates. In this context, it may not be surprising that deISGylase activity is dispensable for IFN sensitivity in human cancer cells. The contribution of deISGylase activity may differ in mice and other species because mouse USP18 efficiently removed ISG15 from substrate proteins (Basters et al., 2014; Ketscher et al., 2015). Species-specific differences in deISGylase activity may reflect evolutionary adaptions to host-pathogen interactions and contribute to divergent viral susceptibility phenotypes observed with *ISG15* loss-of-function mutations (Speer et al., 2016). Although ISG15 is weakly conserved across species, the catalytic site of USP18 is well conserved (Basters et al., 2017; Qiu et al., 2020). This conservation may indicate that the residues required for catalytic function are also required for proper tertiary structure and/or scaffold function(s). Alternatively, human USP18 may have other protein substrates, in addition to conjugated ISG15, for which it is a more efficient enzyme. Indeed, other species, such as birds, do not appear to encode ISG15 in their genomes, although USP18 is conserved (Magor et al., 2013; Qian et al., 2016).

Since ISGylation is tightly coupled to the scaffold role of USP18, caution must be taken when dissecting the catalytic and scaffold roles of USP18. IFN signaling induces the expression of ISGs, including all the machinery required for ISGylation (UBA7, UBCH8, HERC5, ISG15), and ISG15 conjugation occurs concurrently with protein translation (Durfee et al., 2010; Jimenez Fernandez et al., 2019). Therefore, when USP18 scaffold function is lost, more ISGs are synthesized, and these become ISGylated upon translation. Hence, loss of scaffold function results in a large increase of ISGylated proteins, many of which are ISGs themselves (Figure S1). At the same time, loss of deISGylase function can also lead to increased ISGylated proteins. In this study, we designed in vitro assays with purified recombinant protein to directly assess deISGylase activity without the confound of increased ISGylation downstream of impaired scaffold function. Our results demonstrate that I60N retains de-ISGylase activity and the observed accumulation of ISGylated proteins c be decoupled from the loss of catalytic function.

Partial impairments in scaffold function by the I60N mutation correlated with intermediate levels of ISG induction, ISGylation, and IFN sensitivity, implicating scaffold function as the primary mechanism by which USP18 loss promotes tumor intrinsic growth arrest and cell death. Nonetheless, I60N could disrupt additional functions of USP18 that have yet to be described (Martin-Fernandez et al., 2022; Vasou et al., 2021). For example, recent studies have described additional protein-protein interactions with NEMO in the absence of exogenous IFN and with STAT1, OAS3, MX1, and ROBO1 upon IFN stimulation (Singh et al., 2022; Yang et al., 2015). In leukemia-derived THP-1 monocytic cells, nuclear USP18 interacts with IRF9 and STAT2 to regulate non-canonical ISG expression and pyroptosis (Arimoto et al., 2023). It is unknown if these interactions are conserved across cancer cell lines. Future genome wide CRISPR screening strategies paired with endogenous IPs of WT, catalytic, and scaffold mutants may help identify effector pathways and protein-protein interactions mediating USP18-dependent IFN sensitivity across cancer types. A deeper mechanistic understanding will help predict which patients will respond to USP18 inhibition and could explain variable responses across cancer types in syngeneic mouse models (Arimoto et al., 2023; Hong et al., 2014). It will be of great interest to determine whether a universal effector pathway downstream of USP18 is conserved across lineages, or if USP18 can regulate multiple effector pathways that converge on tumor intrinsic growth arrest and cell death.

Understanding the regulation of Type I IFN signaling has profound implications in identifying treatments for a broad range of pathophysiological conditions. Type I IFN signaling is well-established as a critical defense against viral infection and malignant cells (McNab et al., 2015; Snell et al., 2017), and a mediator of immune homeostasis and physiological function (Blaszczyk et al., 2016; Cheon et al., 2014). On the other hand, dysregulation of Type I IFN signaling can result in severe autoimmune disease and has been linked to immunotherapy resistance (Boukhaled et al., 2021; Cheon et al., 2014; Crow and Stetson, 2022; Zitvogel et al., 2015). This work galvanizes our mechanistic understanding of USP18, the key regulator of Type I IFN signaling in human cells, which will facilitate USP18-targeted therapeutics to improve patients’ lives.

## SUPPLEMENTAL INFORMATION

Supplementary Figures S1-3 accompany the paper.

## ACKNOWLEDGMENTS

We thank Jon Oyer and Rubina Tuladhar for generously sharing the CRISPR LAPSE method ahead of publication; Ellene Mashalidis for purification of recombinant SARS-CoV-2 PLpro; Jeffrey Stock for sgRNA and HDR donor oligo design; Kim F. Fennell for recombinant protein efforts; Brooke Conti Trousdale, Jennie Altman, and Drew Nager for comments on the manuscript; Pfizer’s Worldwide Research, Development and Medical Postdoctoral Program for their support; Pfizer colleagues in Emerging Science and Innovation for stimulating discussion; and David Shields for his leadership and thoughtful feedback.

This work was supported by Pfizer and APF and BMK are funded by the Chinese Academy of Medical Sciences (CAMS) Innovation Fund for Medical Science (CIFMS), China (grant nr - 2018-I2M-2-002).

## AUTHOR CONTRIBUTIONS

Experiments were performed by the following authors: HW and VJ (CRISPR LAPSE), VJ (HTRF), HW and CWL (cell line generation and characterization), CWL (endogenous deISGylation), DRH (ISG15-Rh110 deISGylation), KL (UBA7/USP18 double KO characterization), BY and AG (in vivo models), MY (pro-ISG15 cleavage). FW, PDW, VF, VJ, PL, ACG, ECR, BMK, and APF contributed to conception, design, and/or interpretation of the work. VJ, VF, and PDW wrote the manuscript with input from all authors.

## MATERIALS AND METHODS

### CRISPR LAPSE

For *USP18* KO LAPSE 1 × 10^6^ cells were electroporated (Lonza, 4D-Nucleofactor) using the following electroporation buffers and protocols: A549: Buffer SF, CM-130; HAP1: Buffer SE, DZ-113; HCC1143: Buffer SE, E0-117; HCC366: Buffer SE, E0-117; HCT116: Buffer

SE, EN-113; HT-29: Buffer SE, CM-150. For all cell lines, ribonucleoprotein complexes consisting of 200 pmol sgRNA (Thermo Fisher, modified TrueGuide sgRNA) and 10 μg Cas9 (Thermo Fisher, A36498) were prepared in electroporation buffer to a final volume of 100 μL per sample. At 72 h post-electroporation (time initial), cell pools were mixed thoroughly and split into 2 equivalent cell pools maintained in culture for 14 days (time final), with or without 1,000 U/mL Human IFN-Alpha2b (pbl Assay Science, 11105-1). Media was replaced 2x per week and cells were split as needed. At time initial and time final, an aliquot of at least 100,000 cells was pelleted and genomic DNA was extracted (Fisher Scientific, NC9904870) for PCR and Sanger sequencing. Editing efficiency for time initial, time final-IFN, and time final +IFN groups were evaluated using the ICE analysis tool (Synthego). *USP18* KO sgRNA were targeted to the 3’ end of USP18 exon 2, downstream of predicted start codons for both isoforms and upstream of the catalytic cysteine, C64.

#### *USP18* KO

1. sgRNA: GCAGCCCAGAGAGCGTCCCA
2. PCR primers: (Forward, 5’-TCTAAGACCTGCTCTTTGGCATCAGAA-3’; Reverse, 5’-AAAGCTCTGCGGCTTCAGAGCGGAGG-3’)

For USP18 C64S KI CRISPR LAPSE, 1 × 10^6^ cells were electroporated (Maxcyte STx) in 50×3 processing assemblies using electroporation buffer (Maxcyte, EPB-1) and cell-line specific protocols recommended by the manufacturer. Ribonucleoprotein complexes consisting of 200 pmol sgRNA (Synthego, CRISPRevolution sgRNA EZ kit), 10 μg Cas9 (Thermo Fisher, A36499), 4 μM ssODN donor template (IDT) were prepared in electroporation buffer to a final volume of 50 μL per sample. Following electroporation, the transfection mix was equilibrated for 20 min at 37°C. After equilibration, 1 μM HDR Enhancer V2 (IDT, 10007910) was added, and cells were subjected to a cold shock incubation for 48 h at 32°C. At 24 h post-electroporation, media containing 1 μM HDR Enhancer V2 was replaced with fresh media and cells were returned to 32°C until transfer to 37°C at 48 h post-electroporation. At 72 h post-electroporation (time initial), cell pools were mixed thoroughly and split into 2 equivalent cell pools maintained in culture for 14 days (time final), with or without 1,000 U/mL Human IFN-Alpha2b (pbl Assay Science, 11105-1). Media was replaced 2x per week and cells were split as needed. At time initial and time final, an aliquot of 100,000 cells were pelleted and genomic DNA was extracted (Qiagen, 69506) for PCR and Sanger sequencing. Editing efficiency for time initial, time final -IFN, and time final +IFN groups were evaluated using the ICE analysis tool (Synthego).

#### C64S KI

1. sgRNA: ACACCTGAATCAAGGAGTTA
2. ssODN template: CTTATAGGCCTGGTTGGTTTACACAACATTGGACAGACC**AGC**TGTCTTAACTCCTTGAT-TCAGGTGTTCGTAATGAATGTGGACTTCACC
3. PCR primers: (Forward, 5’-CGGCTGATTTTTGGATTTTTTTTGTAGAGACAT-3’; Reverse, 5’-ATAATACAAATGCTCATCAGCCTGAA-3’)

### Purification of recombinant protein

The open reading frame DNA sequences of human USP18(AA16-372) (Uniprot Q9UMW8) and human pro-ISG15(AA1-165)C78S (Uniprot P05161) were synthesized by GENEWIZ. C78S was substituted to stabilize ISG15 for purification (Narasimhan, et al 2005). USP18 was cloned into pFastBac1 between BamHI and XhoI sites with an N-terminal His tag and ISG15 was inserted into pGEX6p1 (Cytiva, 28-9546-48) between BamHI and XhoI sites with an N-terminal GST tag. USP18 C64S, C64S/C65S, and I60N mutations were introduced by quick-change mutagenesis and verified by Sanger sequencing. All USP18 variants were expressed in Spodoptera frugiperda (Sf9, ATCC CRL-1711) insect cell using the Bac-to-Bac Baculovirus Expression System (Invitrogen, 10359016). Sf9 cells were infected at a cell density of 4 × 10^6^cells/mL with a multiplicity of infection of 5.0. At 48 h after infection, cells were collected and resuspended in the lysis buffer containing 25 mM HEPES pH 8.0, 200 mM NaCl, 1 mM TCEP, 40 mM imidazole, EDTA-free complete protease inhibitor cocktail (Sigma, 56619600) and 50 U/ml Pierce universal nuclease (Thermo Fisher, 88702). Sf9 cells were lysed by sonication and His-USP18 protein was purified by Ni-affinity chromatography, His-tag cleavage by PreScission protease and size exclusion chromatography with the SEC buffer 25 mM HEPES pH 8.0, 200 mM NaCl, 1 mM TCEP. GST-ISG15 was expressed in Escherichia coli BL21(DE3) (Thermo Fisher, EC0114) at OD600 0.6-0.8 with 0.1 mM IPTG and 20°C and lysed in the lysis buffer by sonication. After GST-affinity chromatography and tag cleavage by PreScission protease, ISG15 was further purified by size exclusion chromatography (SEC) with the SEC buffer.

The codon-optimized gene corresponding to SARS-CoV-2 PLpro domain (Nsp3 residues 1562-1878; Uniprot P0DTD1) was synthesized as a gBlock gene fragment by Integrated DNA Technologies and was subcloned into a pET28a vector using NcoI and XhoI restriction enzymes (New England Biolabs). The protein was expressed with uncleavable N-terminal Avi and His6x tags in BL21(DE3) E. coli cells (Novagen SinglesTM). The protein was isolated from the lysate using Ni-NTA Agarose resin (Qiagen) and was purified to homogeneity and >95% purity by size exclusion chromatography using a HiLoad Superdex 75 pg 16/600 column (Cytiva) in a buffer containing 50 mM HEPES pH 8, 200 mM NaCl, 1 mM TCEP, and 5% glycerol.

### Genetically modified cell lines

To generate USP18 C64S KI and USP18 I60N KI cells, 1 × 10^6^HAP1 cells were electroporated (Maxcyte STx) in 50×3 processing assemblies using electroporation buffer (Maxcyte, EPB-1) and cell-line specific protocols recommended by the manufacturer. Ribonucleo-protein complexes consisting of 200 pmol sgRNA (Synthego, CRISPRevolution sgRNA EZ kit), 10 μg Cas9 (Thermo Fisher, A36499), 4 μM ssODN donor template (IDT) were prepared in electroporation buffer to a final volume of 50 μL per sample. Following electroporation, the transfection mix was equilibrated for 20 min at 37°C. After equilibration, 1 μM HDR Enhancer V2 (IDT, 10007910) was added and cells were subjected to a cold shock incubation for 48 h at 32°C. At 24 h post-electroporation, media containing 1 μM HDR

Enhancer V2 was replaced with fresh culture media and cells were returned to 32°C until transfer to 37°C at 48 h post-electroporation. Cells were subjected to single-cell cloning on the CellCelector (Sartorius) into 96-well plates. Single-cell clones were confirmed by Sanger sequencing.

#### C64S

1. sgRNA: ACACCTGAATCAAGGAGTTA
2. ssODN template: CTTATAGGCCTGGTTGGTTTACACAACATTGGACAGACC**AGC**TGTCTTAACTCCTTGAT-TCAGGTGTTCGTAATGAATGTGGACTTCACC
3. PCR primers: (Forward, 5’-CGGCTGATTTTTGGATTTTTTTTGTAGAGACAT-3’; Reverse, 5’-ATAATACAAATGCTCATCAGCCTGAA-3’)

#### I60N

1. sgRNA: CTGGTTGGTTTACACAACAT
2. ssODN template: CTTGCCCTCAGCATTTTTTTCTCTTCCCCTTATAGGCCTGGTTGGTTTACACAAT**AAC-** GGACAGACCTGCTGCCTTAACTCCTTGATTCAGGTGTTCGT
3. PCR primers: (Forward, 5’-CGGCTGATTTTTGGATTTTTTTTGTAGAGACAT-3’; Reverse, 5’-AAAGTACCAATACTTGGGTCG-CACCC-3’)

To generate HAP1 *UBA7* KO and *UBA7/USP18* double KO cells in Figure 6, 1 × 10^6^ WT HAP1 cells (Horizon Discovery, 631) or *USP18* KO HAP1 cells (Horizon Discovery, HZGHC000492c006) were electroporated (Maxcyte STx) in OC25×3 processing assemblies using electroporation buffer (Maxcyte, EPB-1) and optimization protocol OP9. Ribonucleoprotein complexes consisting of 200 pmol sgRNA (Synthego, CRISPRevolution sgRNA EZ kit) and 10 μg Cas9 (Thermo Fisher, A36499) were prepared in electroporation buffer to a final volume of 25 μL per sample. Following electroporation, the transfection mix was equilibrated for 20 min at 37°C before addition of fresh media for cell recovery. Knockout efficiency was confirmed by Sanger sequencing and ICE analysis tool (Synthego).

#### UBA7 KO and UBA7/USP18 dKO

1. sgRNA for *UBA7* KO cells: CCCATCCACAGGTATGTGCT
2. sgRNA for *UBA7/USP18* double KO cells: CCCTGAATCCTCTGCATGGC
3. PCR primers: (Forward, 5’-GACCTGGACAGCTCTGTTGA-3’, Reverse, 5’-GCCACCTACACAAAGACCCT-3’)

### Cell viability and qRT-PCR assays

In Figures 6C, 6D, HAP1 cells were plated in 96-well plates (Corning, 3585) at 10,000 cells per well and were subjected to a 9-point dose response of human Interferon Beta 1a (pbl Assay Science, 11410-2) starting at 27,000 U/mL with 3-fold dilution series. Cell viability was measured during IFN treatment (Sartorius, Incucyte S3). After 48 h image collection, cells were lysed for gene expression analysis. Cell lysis, RNA extraction, and cDNA synthesis were performed with the Cells-to-CT gene expression kit (Invitrogen, AM1729). qRT-PCR was performed on the Applied Biosystems QuantStudio 7 Real-Time PCR System according to the manufacturer’s instructions with the following TaqMan gene expression assays: *IFIT1* (Hs0035663_g1), *ISG15* (Hs00192713_m1), and *GAPDH* (Hs99999905_m1). Results shown as expression 2^-ΔCt^ relative to GAPDH. In Figure 5C, qRT-PCR was performed as described above, except that HAP1 cells were plated at 20,000 cells per well.

In Figures 5D, 5E, cells were subjected to an 11-point dose response of Human IFN-alpha2b (pbl Assay Science, 11105-1) starting at 9,000 U/mL with 3.33 dilution series. Cells were plated in 384-well white microplates (Corning, 3570) at 3,000 cells/well in 50 μL media containing IFN-α for 72 h prior to cell lysis with 50 μL CellTiterGlo 2.0 (Promega, G9243). The reaction was incubated in the dark for 20 min at RT. Luminescence was measured on the GloMax Discover (Promega, GM3000) and viability was normalized to 0 U/mL IFN control for each cell line.

### Protein expression assays

1 ×10^5^cells in 24-well plates were treated for 24 h in media containing 1,000 U/mL Human IFN-beta 1a (pbl Assay Science, 11410-2). Lysates were prepared in RIPA buffer (Thermo Scientific, 89900) containing 1x protease and phosphatase inhibitors (Thermo Scientific, 78440) and 250 U/mL benzonase (Sigma, E1014). Concentrations were measured using the Pierce BCA Protein Assay Kit (Thermo Fisher, 23228). Protein analysis was performed on a Wes system (ProteinSimple, 004-600) according to manufacturer’s instructions using a 12-230 kDa Separation Module (ProteinSimple, PS-PP03) and an anti-rabbit Detection Module (ProteinSimple, DM-001). The following primary antibody dilutions were used: rabbit anti-ISG15 (7H29L24) 1:50 (Thermo Fisher, 703131), rabbit anti-USP18 (D4E7) 1:25 (Cell Signaling, 4813S), and rabbit anti-Vinculin 1:300 (Abcam, Ab129002).

### ISG15-Rho110 biochemical assay

Enzymes and ISG15 substrates were diluted 2x in 20 mM Tris-HCl pH 7.5 (Thermo Fisher 15567027), 150 mM NaCl, 0.05% CHAPS (Millipore Sigma C5070), 0.05% BSA (Thermo Fisher 15260037), and 0.5 mM TCEP (UBP Bio CHT#P1021-100). Recombinant human USP18 (WT, C64S, I60N) or mouse WT USP18 were incubated with recombinant human ISG15-Rho110 (South Bay Bio, SBB-PS0002) or mouse ISG15-Rho110 (Ac-ISG15prox-Rho110MP, UbiQ Bio BV, UBIQ-127). Enzymes and substrates were combined at the indicated concentrations in a black 384-well plate (Costar, 3820), briefly centrifuged, and immediately read at an interval of 2 min on an EnVision 2104 reader with excitation filter 485 nm and emission filter 531 nm. For progression curves in Figures S1A, enzymes and substrates were mixed at the indicated concentration and fluorescence from the cleaved substrate was monitored and recorded in the plate reader every two minutes for the indicated time. For Km determinations in Figure 3B, Km/(Km apparent) was calculated in accordance with standard methods. Briefly, a matrix of enzyme concentration ranges was titrated across a range of fluorescent human or mouse ISG15-Rho110 substrate concentrations, and the resultant reactions generated liberated fluorophore from the substrate. The resultant fluorescence was monitored and quantified in a plate reader every two minutes for at least 30 min. From these data, RFU versus time progression curves were generated and initial linearity was determined by non-linear regression in Prism for each enzyme and substrate combination. Enzyme and substrate combinations that resulted in high and linear initial velocities of the reaction were then included in a plot of the velocity (RFUs/time) versus substrate concentration. The velocity versus substrate concentration curve was curve fit in Prism by Michaelis-Menten method for substrate versus velocity to determine Km/(Km apparent) and Vmax/(Vmax apparent). The reported values are an average of two determinations at different enzyme concentrations rounded to two significant digits.

### pSTAT1/total STAT1 HTRF assay

20,000 HAP1 cells were seeded in 96-well plates for 16-24 h with IMDM phenol-red free media (Gibco, 21056023) + 10% HI-FBS (Thermo Fisher, 16140071) prior to adding indicated concentration of human IFN-alpha 2b (pbl Assay Science, 11105-1) for 4 h IFN prime. Cells were washed 1x with PBS and fresh media without IFN was added to allow cells to rest. After 16 – 24 h rest, the indicated concentration of human IFN-alpha 2b (pbl Assay Science, 11105-1) was added for a 15 min pulse prior to cell lysis. pSTAT1 and total STAT1 levels were measured according to manufacturer’s protocol for pSTAT1 Tyr701 (Perkin Elmer, 63ADK026PEG) and total STAT1 (Perkin Elmer, 63ADK096PEG) HTRF kits in HTRF 96-well low volume plates (Perkin Elmer, 66PL96025). HTRF signal was read on an EnVision 2105 reader with excitation filter 320 nm and emission filters 665 nm and 615 nm. pSTAT1/total STAT1 levels were normalized as ratio to account for different IFN concentrations during the 4 h prime.

### In vitro deISGylation of endogenous substrates

HAP1 *USP18* KO cells (Horizon Discovery, HZGHC000492c006) were treated for 24 h with 1,000 U/mL Human IFN-alpha 2b (pbl Assay Science, 11105-1) prior to cell lysis in 20 mM Tris-HCl pH 7.5, 150 mM NaCl, 1% NP40, 0.05% BSA, 0.05% CHAPS, and 0.5 mM TCEP. Lysate concentration was measured using Pierce BCA Protein Assay Kit (Thermo Fisher, 23228), adjusted to 1 μg/μL, and incubated with indicated concentrations of recombinant human USP18 (WT, C64S, I60N), mouse USP18, or SARS-CoV-2 PLpro for 1 h at RT. Western blot analysis was performed on a Wes system (ProteinSimple, 004-600) according to manufacturer’s instructions using a 12-230 kDa Separation Module (ProteinSimple, PS-PP03) and anti-rabbit Detection Module (ProteinSimple, DM-001). The following primary antibody dilutions were used: rabbit anti-ISG15 (7H29L24) 1:50 (Thermo Fisher, 703131), and rabbit anti-Vinculin 1:500 (Abcam, Ab129002).

### Pro-ISG15 cleavage assay

1 μM recombinant human USP18 (WT, C64S, C64S/C65S, I60N) and 5 μM recombinant human pro-ISG15 were incubated for 3 or 10 min at 37°C in the buffer of 25 mM HEPES pH 8.0, 200 mM NaCl, 1 mM TCEP. Cleavage of pro-ISG15(AA1-165) to mature ISG15(AA1-157) was monitored by SDS-PAGE as well as mass spectrometry using 6545 LC/Q-TOF (Agilent technologies). The intact mass analysis was conducted in positive ion mode. The nozzle voltage was 2 kV, capillary voltage was 4 kV, source temperature was 325°C, and the dry gas flow rate was 10 L/min. The protein (10 μL) was loaded by an LC system (Agilent 1290 Infinity II, Agilent Technologies, Santa Clara, CA) on a HPLC column (PLRP-S 1000 Å, 5 mm, 50×2.1 mm, Agilent Technologies, Santa Clara, CA). The following solvents were used: solvent A, water with 0.1% formic acid; solvent B, acetonitrile with 0.1% formic acid. The protein was eluted at 400 μL/min with the following gradient: 2% B in 1 min, 85% B in 9 min, 95% B in 0.1 min, 95% B from 10.1-12.0 min then went back to 2% in 0.1 min. For data analysis, spectra were coadded and deconvoluted using Agilent MassHunter software with a 0.5 Da mass step and mass range of 10-150 kDa.

### Syngeneic mouse models

Two rounds of electroporation were performed to generate *Usp18* KO CT26 cells. First, WT CT26 cells were electroporated with sgRNA-*Usp18*#1 (TGTACAGCCCACGCAAATC) and Cas9 and expanded for 4 days prior to a second round of electroporation with sgRNA-*Usp18*#2 (AAAGTCAGCCATCCCAACGT) and Cas9. *Rosa26* KO cells were generated by a single round of electroporation with sgRNA-*Rosa26* (GAAGATGGGCGGGAGTCTTC) and Cas9. PCR amplicons surrounding *Usp18* sgRNA cut sites were sequenced (MiSeq, Illumina) and frame-shift mutation frequency for triploid CT26 cells was estimated to be ∼65% by the following equation: [1-((1

– FSsg1) ^*^ (1 – FSsg2))]^3^ where FS = % frame shift mutation for a given sgRNA. Loss-of-function KO frequency may be higher given that large in-frame mutations may also result in loss-of-function.

All procedures performed on animals were in accordance with regulations and established guidelines and were reviewed and approved by an Institutional Animal Care and Use Committee or through an ethical review process. BALB/c mice (strain code – 028), 6-8 weeks old were procured from Charles River Laboratories and housed under standard 12:12 light:dark cycle in ventilated racks at a room temp of 72ºF and RH between 30-70%. After acclimatization, 1 × 10^6^ cells CT26 cells (ATCC, CRL-2638TM) (*Rosa26* or *Usp18* KO) were implanted subcutaneously in the right flank of each mouse. Tumor volume was measured at least twice weekly with a caliper using the longest dimension (length) and the longest perpendicular dimension (width) from Day 6 to Day 27 post-implantation. Tumor volume was estimated with the formula: (L×W^2^)/2.

## SUPPLEMENTARY FIGURES

**Figure S1:**
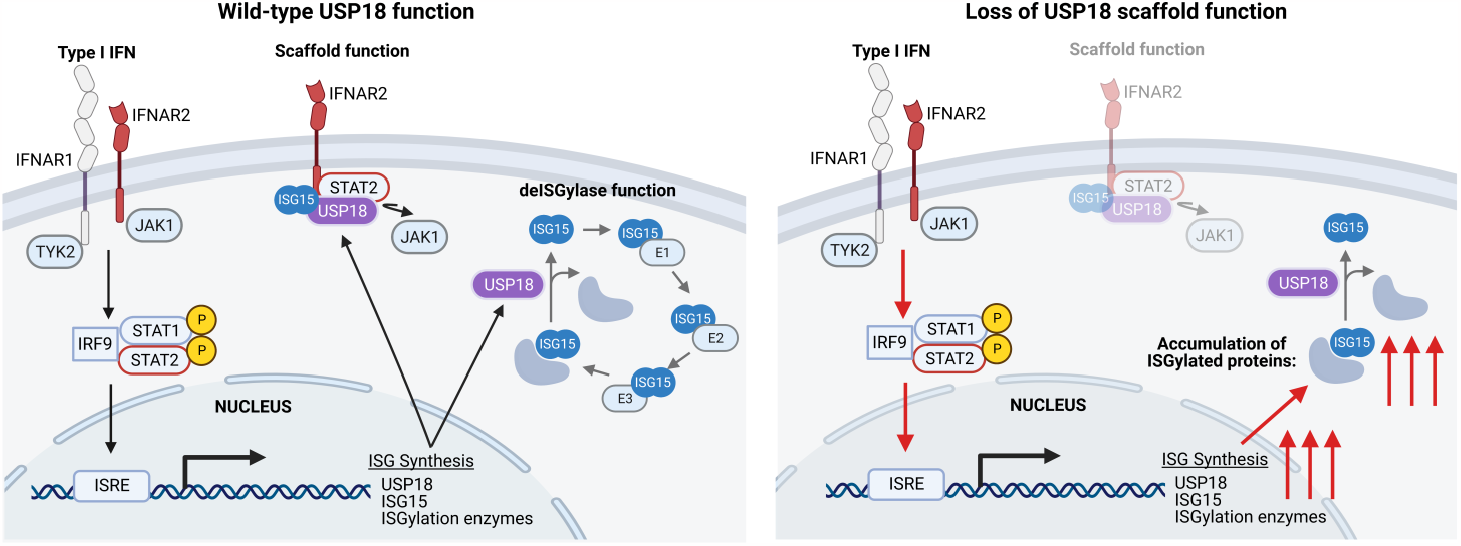
Loss of USP18 scaffold function can increase ISGylated protein pool. Type I IFN activation of the IFNAR signaling cascade results in ISG synthesis. Type I IFN-inducible ISGs include USP18, ISG15, and the enzymes required for ISGylation (UBA7, UBCH8, and HERC5). USP18 negatively regulates IFN responses by displacing JAK1 to repress IFNAR signaling and removing ISG15 from ISGylated proteins (left schematic). Loss of USP18 scaffold function derepresses IFNAR signaling, resulting in hypersensitivity to Type I IFNs, increased ISG synthesis, and increased ISGylation of newly synthesized proteins (right schematic) Schematics created with BioRender.com.

**Figure S2:**
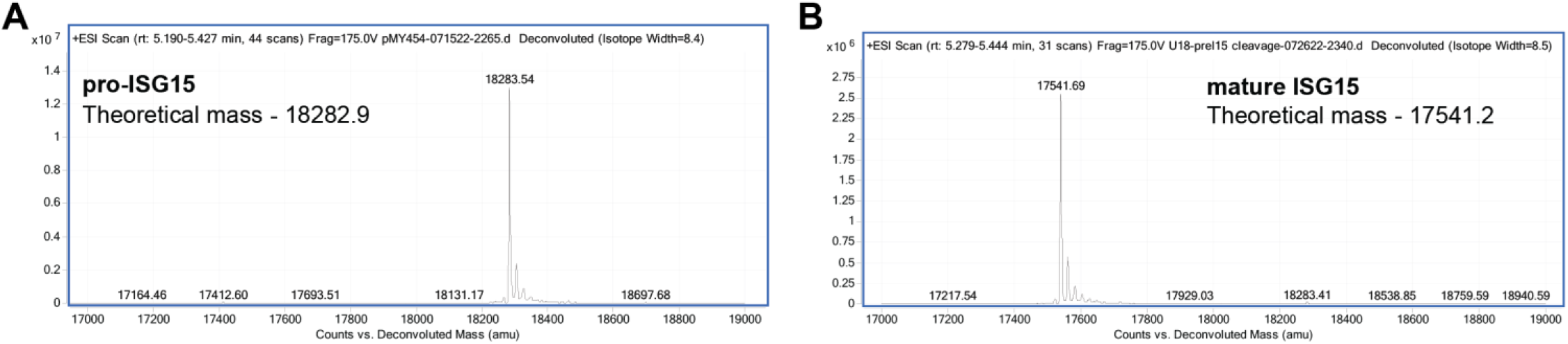
USP18 cleaves pro-ISG15 to produce mature ISG15. (A,B) Cleavage of 5 μM human pro-ISG15 (AA1-165) to mature ISG15 (AA1-157) was assessed by mass spectrometry after 10 min incubation at 37°C with 1 μM of recombinant human WT USP18.

**Figure S3:**
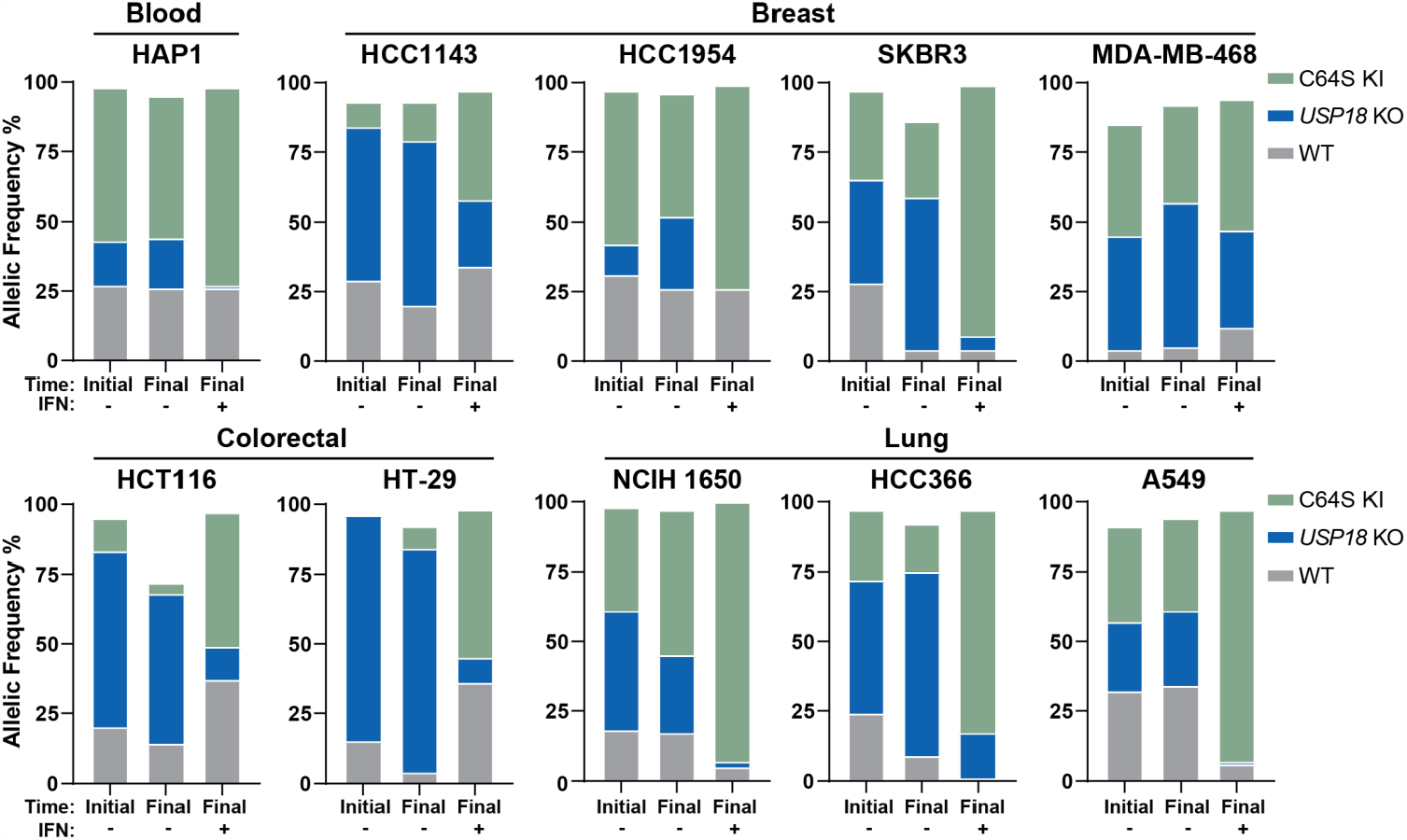
Cancer cells expressing USP18 C64S are not sensitive to IFN. Independent biological replicate run in parallel with Figure 7: parental human cancer cell lines were electroporated with C64S donor oligo and sgRNA targeted to *USP18*. Allelic frequency was determined by sequencing samples 72 h post-electroporation (time initial) and after 2 additional weeks of passaging cells (time final) in the presence (+) or absence (−) of 1000 U/mL IFN-α. KO indicates frame-shift mutation or in-frame mutation ≥ 21 bp; WT indicates no mutation or in-frame mutation < 21 bp; KI indicates donor oligo integration at the intended position.

## REFERENCES

Alsohime, F., Martin-Fernandez, M., Temsah, M.H., Alabdulhafid, M., Le Voyer, T., Alghamdi, M., Qiu, X., Alotaibi, N., Alkahtani, A., Buta, S., et al. (2020). JAK Inhibitor Therapy in a Child with Inherited USP18 Deficiency. N Engl J Med 382, 256–265.

Arimoto, K.I., Lochte, S., Stoner, S.A., Burkart, C., Zhang, Y., Miyauchi, S., Wilmes, S., Fan, J.B., Heinisch, J.J., Li, Z., et al. (2017). STAT2 is an essential adaptor in USP18-mediated suppression of type I interferon signaling. Nat Struct Mol Biol 24, 279–289.

Arimoto, K.I., Miyauchi, S., Troutman, T.D., Zhang, Y., Liu, M., Stoner, S.A., Davis, A.G., Fan, J.B., Huang, Y.J., Yan, M., et al. (2023). Expansion of interferon inducible gene pool via USP18 inhibition promotes cancer cell pyroptosis. Nat Commun 14, 251.

Barber, D.L., Wherry, E.J., Masopust, D., Zhu, B., Allison, J.P., Sharpe, A.H., Freeman, G.J., and Ahmed, R. (2006). Restoring function in exhausted CD8 T cells during chronic viral infection. Nature 439, 682–687.

Basters, A., Geurink, P.P., El Oualid, F., Ketscher, L., Casutt, M.S., Krause, E., Ovaa, H., Knobeloch, K.P., and Fritz, G. (2014). Molecular characterization of ubiquitin-specific protease 18 reveals substrate specificity for interferon-stimulated gene 15. FEBS J 281, 1918–1928.

Basters, A., Geurink, P.P., Rocker, A., Witting, K.F., Tadayon, R., Hess, S., Semrau, M.S., Storici, P., Ovaa, H., Knobeloch, K.P., et al. (2017). Structural basis of the specificity of USP18 toward ISG15. Nat Struct Mol Biol 24, 270–278.

Bekisz, J., Baron, S., Balinsky, C., Morrow, A., and Zoon, K.C. (2010). Antiproliferative Properties of Type I and Type II Interferon. Pharmaceuticals (Basel) 3, 994–1015.

Blaszczyk, K., Nowicka, H., Kostyrko, K., Antonczyk, A., Wesoly, J., and Bluyssen, H.A. (2016). The unique role of STAT2 in constitutive and IFN-induced transcription and antiviral responses. Cytokine Growth Factor Rev 29, 71–81.

Boukhaled, G.M., Harding, S., and Brooks, D.G. (2021). Opposing Roles of Type I Interferons in Cancer Immunity. Annu Rev Pathol 16, 167–198.

Cheon, H., Borden, E.C., and Stark, G.R. (2014). Interferons and their stimulated genes in the tumor microenvironment. Semin Oncol 41, 156–173.

Corey, D.R., and Craik, C.S. (1992). An Investigation into the Minimum Requirements for Peptide Hydrolysis by Mutation of the Catalytic Triad of Trypsin. J Am Chem Soc 114, 1784–1790.

Craik, C.S., Roczniak, S., Largman, C., and Rutter, W.J. (1987). The Catalytic Role of the Active Site Aspartic Acid in Serine Proteases. Science 237, 909–913.

Crow, Y.J., and Stetson, D.B. (2022). The type I interferonopathies: 10 years on. Nat Rev Immunol 22, 471–483.

Desai, S.D., Haas, A.L., Wood, L.M., Tsai, Y.C., Pestka, S., Rubin, E.H., Saleem, A., Nur, E.K.A., and Liu, L.F. (2006). Elevated expression of ISG15 in tumor cells interferes with the ubiquitin/26S proteasome pathway. Cancer Res 66, 921–928.

Duncan, C.J.A., Thompson, B.J., Chen, R., Rice, G.I., Gothe, F., Young, D.F., Lovell, S.C., Shuttleworth, V.G., Brocklebank, V., Corner, B., et al. (2019). Severe type I interferonopathy and unrestrained interferon signaling due to a homozygous germline mutation in STAT2. Science Immunology 4.

Durfee, L.A., Lyon, N., Seo, K., and Huibregtse, J.M. (2010). The ISG15 conjugation system broadly targets newly synthesized proteins: implications for the antiviral function of ISG15. Mol Cell 38, 722–732.

Fan, J.B., Arimoto, K., Motamedchaboki, K., Yan, M., Wolf, D.A., and Zhang, D.E. (2015). Identification and characterization of a novel ISG15-ubiquitin mixed chain and its role in regulating protein homeostasis. Sci Rep 5, 12704.

Ferreira, J.C., Fadl, S., and Rabeh, W.M. (2022). Key dimer interface residues impact the catalytic activity of 3CLpro, the main protease of SARS-CoV-2. J Biol Chem 298, 102023.

Francois-Newton, V., Magno de Freitas Almeida, G., Payelle-Brogard, B., Monneron, D., Pichard-Garcia, L., Piehler, J., Pellegrini, S., and Uze, G. (2011). USP18-based negative feedback control is induced by type I and type III interferons and specifically inactivates interferon alpha response. PLoS One 6, e22200.

Freitas, B.T., Durie, I.A., Murray, J., Longo, J.E., Miller, H.C., Crich, D., Hogan, R.J., Tripp, R.A., and Pegan, S.D. (2020). Characterization and Noncovalent Inhibition of the Deubiquitinase and deISGylase Activity of SARS-CoV-2 Papain-Like Protease. ACS Infect Dis 6, 2099–2109.

Gessani, S., Conti, L., Del Corno, M., and Belardelli, F. (2014). Type I interferons as regulators of human antigen presenting cell functions. Toxins (Basel) 6, 1696–1723.

Giannakopoulos, N.V., Arutyunova, E., Lai, C., Lenschow, D.J., Haas, A.L., and Virgin, H.W. (2009). ISG15 Arg151 and the ISG15-conjugating enzyme UbE1L are important for innate immune control of Sindbis virus. J Virol 83, 1602–1610.

Gruber, C., Martin-Fernandez, M., Ailal, F., Qiu, X., Taft, J., Altman, J., Rosain, J., Buta, S., Bousfiha, A., Casanova, J.L., et al. (2020). Homozygous STAT2 gain-of-function mutation by loss of USP18 activity in a patient with type I interferonopathy. J Exp Med 217.

Guo, Y., Chinyengetere, F., Dolinko, A.V., Lopez-Aguiar, A., Lu, Y., Galimberti, F., Ma, T., Feng, Q., Sekula, D., Freemantle, S.J., et al. (2012). Evidence for the ubiquitin protease UBP43 as an antineoplastic target. Mol Cancer Ther 11, 1968–1977.

Hong, B., Li, H., Lu, Y., Zhang, M., Zheng, Y., Qian, J., and Yi, Q. (2014). USP18 is crucial for IFN-γ-mediated inhibition of B16 melanoma tumorigenesis and antitumor immunity. Molecular Cancer 13.

Jimenez Fernandez, D., Hess, S., and Knobeloch, K.P. (2019). Strategies to Target ISG15 and USP18 Toward Therapeutic Applications. Front Chem 7, 923.

Kang, D., Jiang, H., Wu, Q., Pestka, S., and Fisher, P.B. (2001). Cloning and characterization of human ubiquitin-processing protease-43 from terminally differentiated human melanoma cells using a rapid subtraction hybridization protocol RaSH. Gene 267, 233–242.

Ketscher, L., Hannss, R., Morales, D.J., Basters, A., Guerra, S., Goldmann, T., Hausmann, A., Prinz, M., Naumann, R., Pekosz, A., et al. (2015). Selective inactivation of USP18 isopeptidase activity in vivo enhances ISG15 conjugation and viral resistance. Proc Natl Acad Sci U S A 112, 1577–1582.

Lai, C., Struckhoff, J.J., Schneider, J., Martinez-Sobrido, L., Wolff, T., Garcia-Sastre, A., Zhang, D.E., and Lenschow, D.J. (2009). Mice lacking the ISG15 E1 enzyme UbE1L demonstrate increased susceptibility to both mouse-adapted and non-mouse-adapted influenza B virus infection. J Virol 83, 1147–1151.

Magor, K.E., Miranzo Navarro, D., Barber, M.R., Petkau, K., Fleming-Canepa, X., Blyth, G.A., and Blaine, A.H. (2013). Defense genes missing from the flight division. Dev Comp Immunol 41, 377–388.

Malakhov, M.P., Kim, K.I., Malakhova, O.A., Jacobs, B.S., Borden, E.C., and Zhang, D.E. (2003). High-throughput immunoblotting. Ubiquitiin-like protein ISG15 modifies key regulators of signal transduction. J Biol Chem 278, 16608–16613.

Malakhov, M.P., Malakhova, O.A., Kim, K.I., Ritchie, K.J., and Zhang, D.E. (2002). UBP43 (USP18) specifically removes ISG15 from conjugated proteins. J Biol Chem 277, 9976–9981.

Malakhova, O.A., Yan, M., Malakhov, M.P., Yuan, Y., Ritchie, K.J., Kim, K.I., Peterson, L.F., Shuai, K., and Zhang, D.E. (2003). Protein ISGylation modulates the JAK-STAT signaling pathway. Genes Dev 17, 455–460.

Martin-Fernandez, M., Buta, S., Le Voyer, T., Li, Z., Dynesen, L.T., Vuillier, F., Franklin, L., Ailal, F., Muglia Amancio, A., Malle, L., et al. (2022). A partial form of inherited human USP18 deficiency underlies infection and inflammation. J Exp Med 219.

McNab, F., Mayer-Barber, K., Sher, A., Wack, A., and O’Garra, A. (2015). Type I interferons in infectious disease. Nat Rev Immunol 15, 87–103.

Meuwissen, M.E., Schot, R., Buta, S., Oudesluijs, G., Tinschert, S., Speer, S.D., Li, Z., van Unen, L., Heijsman, D., Goldmann, T., et al. (2016). Human USP18 deficiency underlies type 1 interferonopathy leading to severe pseudo-TORCH syndrome. J Exp Med 213, 1163–1174.

Munnur, D., Teo, Q., Eggermont, D., Lee, H.H.Y., Thery, F., Ho, J., van Leur, S.W., Ng, W.W.S., Siu, L.Y.L., Beling, A., et al. (2021). Altered ISGylation drives aberrant macrophage-dependent immune responses during SARS-CoV-2 infection. Nat Immunol 22, 1416–1427.

Mustachio, L.M., Kawakami, M., lu, Y., Rodriguez-Canales, J., Mino, B., Behrens, C., Wistuba, I., Bota-Rabassedas, N., Yu, J., Jack Lee, J., et al. (2017a). The ISG15-specific protease USP18 regulates stability of PTEN. Oncotarget 8, 3–14.

Mustachio, L.M., Lu, Y., Tafe, L.J., Memoli, V., Rodriguez-Canales, J., Mino, B., Villalobos, P.A., Wistuba, I., Katayama, H., Hanash, S.M., et al. (2017b). Deubiquitinase USP18 Loss Mislocalizes and Destabilizes KRAS in Lung Cancer. Mol Cancer Res 15, 905–914.

Okumura, F., Okumura, A.J., Uematsu, K., Hatakeyama, S., Zhang, D.E., and Kamura, T. (2013). Activation of doublestranded RNA-activated protein kinase (PKR) by interferonstimulated gene 15 (ISG15) modification down-regulates protein translation. J Biol Chem 288, 2839–2847.

Park, J.H., Yang, S.W., Park, J.M., Ka, S.H., Kim, J.H., Kong, Y.Y., Jeon, Y.J., Seol, J.H., and Chung, C.H. (2016). Positive feedback regulation of p53 transactivity by DNA damage-induced ISG15 modification. Nat Commun 7, 12513.

Perng, Y.C., and Lenschow, D.J. (2018). ISG15 in antiviral immunity and beyond. Nat Rev Microbiol 16, 423–439.

Pinto-Fernandez, A., Salio, M., Partridge, T., Chen, J., Vere, G., Greenwood, H., Olie, C.S., Damianou, A., Scott, H.C., Pegg, H.J., et al. (2020). Deletion of the deISGylating enzyme USP18 enhances tumour cell antigenicity and radiosensitivity. Br J Cancer.

Potu, H., Sgorbissa, A., and Brancolini, C. (2010). Identification of USP18 as an important regulator of the susceptibility to IFN-alpha and drug-induced apoptosis. Cancer Res 70, 655–665.

Qian, W., Wei, X., Zhou, H., and Jin, M. (2016). Molecular cloning and functional analysis of duck ubiquitin-specific protease 18 (USP18) gene. Dev Comp Immunol 62, 39–47.

Qiu, X., Taft, J., and Bogunovic, D. (2020). Developing Broad-Spectrum Antivirals Using Porcine and Rhesus Macaque Models. J Infect Dis 221, 890–894.

Schneider, W.M., Chevillotte, M.D., and Rice, C.M. (2014). Interferon-stimulated genes: a complex web of host defenses. Annu Rev Immunol 32, 513–545.

Singh, A., Padariya, M., Faktor, J., Kote, S., Mikac, S., Dziadosz, A., Lam, T.W., Brydon, J., Wear, M.A., Ball, K.L., et al. (2022). Identification of novel interferon responsive protein partners of human leukocyte antigen A (HLA-A) using cross-linking mass spectrometry (CLMS) approach. Sci Rep 12, 19422.

Snell, L.M., McGaha, T.L., and Brooks, D.G. (2017). Type I Interferon in Chronic Virus Infection and Cancer. Trends Immunol 38, 542–557.

Speer, S.D., Li, Z., Buta, S., Payelle-Brogard, B., Qian, L., Vigant, F., Rubino, E., Gardner, T.J., Wedeking, T., Hermann, M., et al. (2016). ISG15 deficiency and increased viral resistance in humans but not mice. Nat Commun 7, 11496.

Sprang, S., Standing, T., Fletterick, R.J., Stroud, R.M., Finer-Mooer, J., Xuong, N.-H., Hamlin, R., Rutter, W.J., and Craik, C.S. (1987). The Three-Dimensional Structure of Asn102 Mutant of Trypsin: Role of Asp102 in Serine Protease Catalysis. Science 237, 905–909.

Taft, J., and Bogunovic, D. (2018). The Goldilocks Zone of Type I IFNs: Lessons from Human Genetics. J Immunol 201, 3479–3485.

Tang, Y., Zhong, G., Zhu, L., Liu, X., Shan, Y., Feng, H., Bu, Z., Chen, H., and Wang, C. (2010). Herc5 attenuates influenza A virus by catalyzing ISGylation of viral NEV1 protein. J Immunol 184, 5777–5790.

Vasou, A., Nightingale, K., Cetkovská, V., Bamford, C.G.G., Andrejeva, J., Randall, R.E., McLauchlan, J., Weekes, M.P., and Hughes, D.J. (2021). A co-opted ISG15-USP18 binding mechanism normally reserved for deISGylation controls type I IFN signalling. bioRxiv.

Yang, Z., Xian, H., Hu, J., Tian, S., Qin, Y., Wang, R.F., and Cui, J. (2015). USP18 negatively regulates NF-kappaB signaling by targeting TAK1 and NEMO for deubiquitination through distinct mechanisms. Sci Rep 5, 12738.

Zhang, X., Bogunovic, D., Payelle-Brogard, B., Francois-Newton, V., Speer, S.D., Yuan, C., Volpi, S., Li, Z., Sanal, O., Mansouri, D., et al. (2015). Human intracellular ISG15 prevents interferonalpha/beta over-amplification and auto-inflammation. Nature 517, 89–93.

Zhang, Y., Thery, F., Wu, N.C., Luhmann, E.K., Dussurget, O., Foecke, M., Bredow, C., Jimenez-Fernandez, D., Leandro, K., Beling, A., et al. (2019). The in vivo ISGylome links ISG15 to metabolic pathways and autophagy upon Listeria monocytogenes infection. Nat Commun 10, 5383.

Zhao, C., Hsiang, T.Y., Kuo, R.L., and Krug, R.M. (2010). ISG15 conjugation system targets the viral NEV1 protein in influenza A virus-infected cells. Proc Natl Acad Sci U S A 107, 2253–2258.

Zitvogel, L., Galluzzi, L., Kepp, O., Smyth, M.J., and Kroemer, G. (2015). Type I interferons in anticancer immunity. Nat Rev Immunol 15, 405–414.

